# All-in-one AAV-delivered epigenome-editing platform: *proof-of-concept* and therapeutic implications for neurodegenerative disorders

**DOI:** 10.1101/2023.04.14.536951

**Authors:** Boris Kantor, Bernadette Odonovan, Joseph Rittiner, Dellila Hodgson, Nicholas Lindner, Sophia Guerrero, Wendy Dong, Austin Zhang, Ornit Chiba-Falek

## Abstract

Safely and efficiently controlling gene expression is a long-standing goal of biomedical research, and the recently discovered bacterial CRISPR/Cas system can be harnessed to create powerful tools for epigenetic editing. Current state-of-the-art systems consist of a deactivated-Cas9 nuclease (dCas9) fused to one of several epigenetic effector motifs/domains, along with a guide RNA (gRNA) which defines the genomic target. Such systems have been used to safely and effectively silence or activate a specific gene target under a variety of circumstances. Adeno-associated vectors (AAVs) are the therapeutic platform of choice for the delivery of genetic cargo; however, their small packaging capacity is not suitable for delivery of large constructs, which includes most CRISPR/dCas9-effector systems. To circumvent this, many AAV-based CRISPR/Cas tools are delivered in two pieces, from two separate viral cassettes. However, this approach requires higher viral payloads and usually is less efficient. Here we develop a compact dCas9-based repressor system packaged within a single, optimized AAV vector. The system uses a smaller dCas9 variant derived from *Staphylococcus aureus* (*Sa*). A novel repressor was engineered by fusing the small transcription repression domain (TRD) from MeCP2 with the KRAB repression domain. The final d*Sa*Cas9-KRAB-MeCP2(TRD) construct can be efficiently packaged, along with its associated gRNA, into AAV particles. Using reporter assays, we demonstrate that the platform is capable of robustly and sustainably repressing the expression of multiple genes-of-interest, both *in vitro* and *in vivo*. Moreover, we successfully reduced the expression of ApoE, the stronger genetic risk factor for late onset Alzheimer’s disease (LOAD). This new platform will broaden the CRISPR/dCas9 toolset available for transcriptional manipulation of gene expression in research and therapeutic settings.

## Introduction

Bacterial Clustered Regularly Interspaced Palindromic Repeats (CRISPR)/Cas systems have evolved to bind and cleave nucleic acids in a highly-efficient and flexible fashion [1], [2], [3]. Since their discovery, various Cas orthologs and variants with useful properties have been identified and harnessed for use in gene editing. However, gene-editing approaches employing endonuclease active, wild-type Cas9 enzymes have resulted in unwanted off-target effects, including cell cycle arrest, changes in cellular differentiation, and apoptotic signaling, which are serious barriers against the use of this technology in gene therapy applications [4], [5], [6]. Earlier studies have shown that point mutations introduced into the catalytic domains of Cas9 enzymes can completely abolish their endonuclease activity without affecting their affinity for the targeted DNA [7], [8], [9], [10]. Furthermore, by fusing epigenetic effectors or domains with the catalytically-inactive, or ‘dead’, Cas9 (dCas9), the expression of target genes can be increased or decreased through epigenetic editing [9], [11], [12], [13], [14]. These approaches (repurposing the CRISPR/Cas system for gene activation or repression) have been coined CRISPR-activation (CRISPR*a*) and CRISPR-interference (CRISPR*i*), respectively. A variety of these tools, including those based on DNA methyltransferases (DNMTs), transcriptional repressors, and histone-modifying enzymes (HMEs) have been developed, with many achieving strong levels of gene repression (reviewed in [15]). These approaches are also applicable in a clinical setting, and targeted regulation of disease-causal genes offers novel avenues for the development of a new generation of gene therapies [16], [17], [18]. For example, our laboratory recently developed an all-in-one lentiviral vector expressing dCas9 fused to the catalytic domain of DNA methyltransferase 3A (DNMT3A) for targeted repression of the *SNCA* gene, as a therapeutic strategy for Parkinson’s Disease (PD) [19, 20]. We showed that the reduction in *SNCA* levels mediated by the [*SNCA* gRNA]/dCas9-DMNT3A system successfully rescued disease-related cellular phenotypes including the production of mitochondrial reactive oxygen species (ROS) and cellular viability of dopaminergic neurons [19], [20].

As with many other gene therapy tools, CRISPR/Cas components are commonly delivered *via* viral vectors (reviewed in [15]). Lentiviral vectors (LVs) and adeno-associated vectors (AAVs) offer an effective method for gene-to-cell transfer, and as such these platforms occupy a central place among delivery systems used for gene therapy applications (reviewed in [15]). LVs are attractive delivery vehicles due to their ability to accommodate large transgenic payloads and sustain a robust level of gene expression in a wide range of dividing and non-dividing cells (reviewed in [21]). However, long-term expression of LV-delivered Cas9/guide RNA systems may lead to substantial undesirable off-target perturbations characterized by non-specific RNA-DNA interactions and off-target DNA cleavage [22]. Furthermore, as an integrating system, LVs possess a significant risk of insertional mutagenicity and even oncogenicity [23]. Recombinant AAV vectors (rAAVs), on the other hand, offer a transiently expressing platform, along with very weak integration capacity. Recent advances in the development of preclinically and clinically graded AAVs have propelled this system into broad use in the gene therapy field. Indeed, preclinical and therapeutic successes in AAV-based gene replacement and gene editing have helped AAV to gain a reputation as a leading therapeutic platform, with three AAV-mediated gene therapy products recently gaining regulatory approval in Europe and the United States (reviewed in [24]).

However, a major limitation of using AAV vectors is their relatively small transgene capacity (up to approximately 4.7kb), which makes it difficult (or outright impossible) to package bulky transgenes. For instance, the coding sequence of *Streptococcus pyogenes* Cas9 (SpCas9) is 4.2kb, which consumes nearly all of the packaging room of the vector. The discovery of smaller Cas9 enzymes, including those derived from *Staphylococcus aureus* (SaCas9), *Campylobacter jejuni* (CjCas9), Deltaproteobacteria (CasX), and most recently a miniature Cas system (CasMINI) engineered from the type V-F Cas12f (Cas14), has led to the development of Cas/guide RNA systems which are more suitable to be packaged into AAV vectors [25], [26], [27], [28]. Furthermore, all the above endonucleases have been successfully converted into their respective non-active versions to support various gene-repurposing applications. For example, Thakore and colleagues developed a dSaCas9-KRAB repressor system packaged in AAV particles [29]. Nevertheless, the authors utilized a dual-AAV system, delivering dSaCas9-KRAB and a Pcsk9-targeting gRNA from two separate expression cassettes to repress the transcription of Pcsk9 (a regulator of cholesterol levels) in the liver of adult mice [29]. The value of an improved Cas-effector pairing would be immense, considering that in addition to the CRISPR/Cas components the vector has to accommodate at least two promoters (a traditional Pol II promoter to express the Cas9-repressor protein, and a Pol III promoter to express the gRNA component), a poly(A) signal for transcriptional termination, and nuclear localization signals (NLSs), as well as other *cis-acting* elements such as Woodchuck hepatitis virus Posttranscriptional Regulatory Element (WPRE).

Furthermore, AAV-based systems are significantly more sensitive than other vectors (*e.g.* LVs) when it comes to packaging of multipart components, such as CRISPR/Cas [24]. As a consequence, most of the all-in-one CRISPR/Cas9 tools developed so far are based on plasmid-based or lentiviral delivery systems [30], [31], [32], [33]. We recently demonstrated that the packaging efficiency and viral titers of vector systems bearing large gene-editing tools could be significantly improved by the optimizations made within the vector backbone. Using integrase-deficient (IDLV) and integrase-competent (ICLV) lentiviral vectors, we demonstrated that IDLV-CRISPR/Cas and ICLV-CRISPR/Cas constructs carrying multiple binding sites for the transcription factors Sp1 and NF-kB could be packaged more efficiently and produced at higher titers [22]. Furthermore, functional titers (measured in the transduced cells) also showed a significant improvement compared to their naïve viral counterparts [22]. The improved vectors were able to mediate efficient and robust gene editing perturbations *in vitro* and *in vivo.* Lastly, the IDLV-CRISPR/Cas vector showed only minimal off-target effects, and majority of its genome remains in an episomal (non-integrated) state [22].

Notably, most recombinant vectors, including LVs and AAVs, are lacking many of their endogenous elements, including the above Sp1 and NF-kB sites, which are deleted from vector cassettes along with larger elements, primarily due to safety reasons [19], [22], [34]. Thus, most episomal vectors, including IDLV, AAV, Cytomegalovirus (CMV), Epstein-Barr virus (EBV), and Herpes simplex virus type 1 (HSV-1) inherit the limitation of being epigenetically silenced by default. We and other research groups have studied the early stages of the viral life cycle and reported a competition between cellular epigenetic silencing of viral genes and viral inhibition of repressive factors, such as virus-encoded histone deacetylases (HDACs) and other factors [22], [35], [36], [37], [38]. Therefore, a lack of viral activation machinery recruiting these transcriptional factors could distort the transcriptional environment on the chromatin level, resulting in transcriptional silencing and general impairment of viral expression [22], [39]. AAV is a prominent example as its genome is organized into repressive, silencing chromatin structures [21], [22]. And, as noted above, the expression cassettes of rAAVs are scrubbed of Sp1 and NF-kB binding sites [40], [41], [42]. To address this, we developed and validated an improved AAV vector carrying a concatemer of the Sp1 and NF-kB recognition sites in its expression cassette. We then used the optimized vector backbone to screen for epigenetic editors efficiently packaged into all-in-one AAV particles. The lead platform harbored dSaCas9 fused to transcriptional repression domains (TRDs) domains derived from MeCP2 and KRAB. The system could be efficiently packaged into AAV, and robustly suppressed gene expression *in vitro* and *in vivo*. This novel platform can expand the AAV/CRISPR-dCas9 toolbox for both basic research and preclinical/clinical studies.

## Results

### Optimization of AAV vector backbone

The genome of recombinant AAV harbors no Sp1 and NF-kB binding sites, unlike the wild type virus [19], [22], [34], [40], [41], [42]. As a first step to build an efficient epigenome-editing system for AAV delivery, we decided to reintroduce the above transcription activator binding sites into the expression cassette of the vector. We inserted 2xSp1, 4xSp1, 2xNF-kB, 4xNF-kB, and 2xSp1+2xNF-kB binding sites into the backbone of an AAV vector expressing a destabilized GFP and Nano-Luciferase dGFP/NLuc reporter **(Fig. 1a).** All sites were cloned upstream from the core (minimal) portion of the EF1α promoter, dubbed EFS-NC. The naïve EFS-NC promoter (which carries neither Sp1 nor NF-kB binding sites) was used as a control. In addition, a full-length version of the EF1α promoter, which contains multiple Sp1 and NF-kB sites was used as a positive control. The full-length EF1α promoter is ubiquitous and strong, but its large size (∼1500bps) generally is not suitable for AAV packaging.

**Fig. 1.**
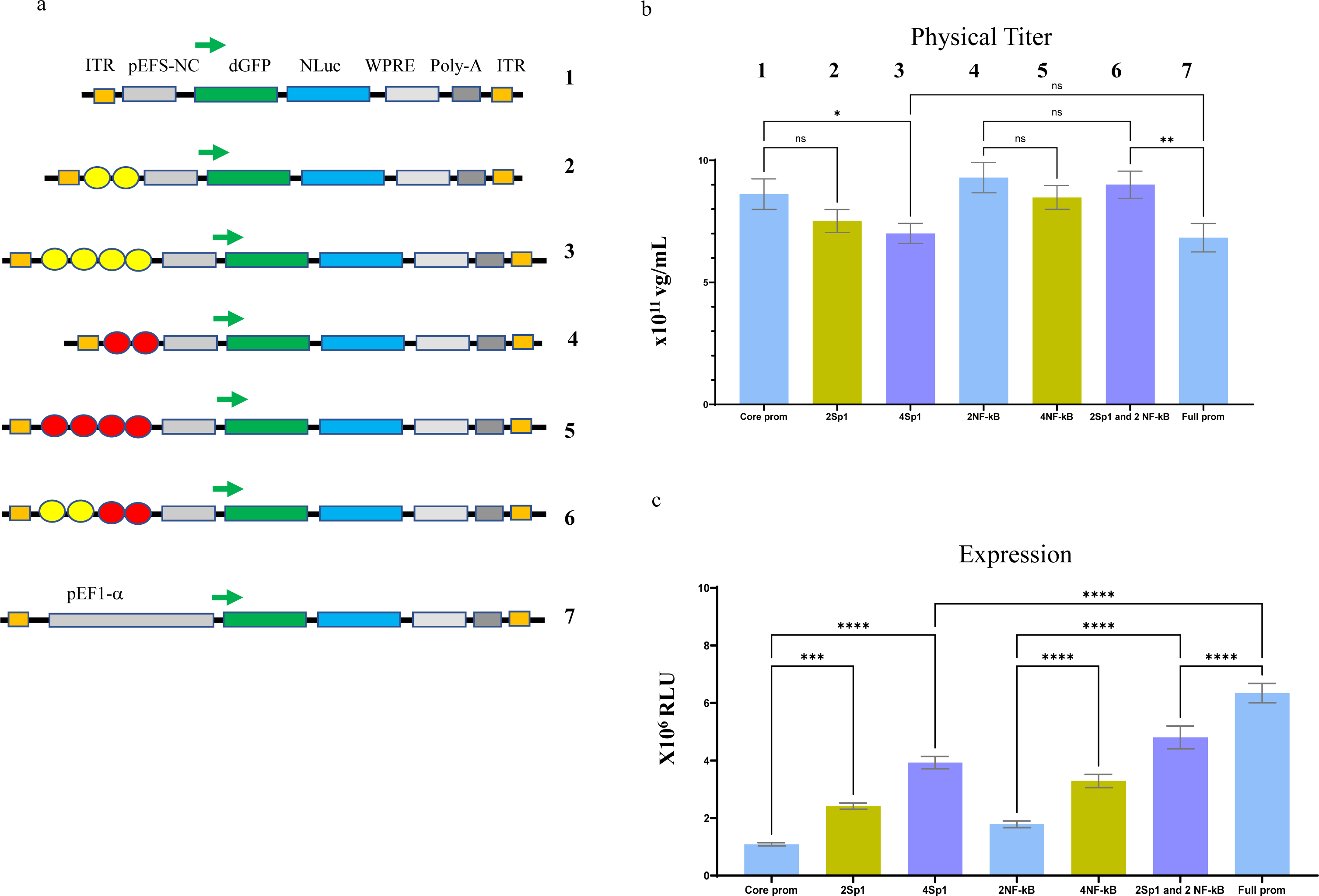

However, the miniature EFS-NC promoter (only 212 bps long) fits most transgenes expressed via AAV systems. To test whether the above modifications would improve the packaging efficiency and transcription, the vectors were first manufactured at non-concentrated grades and titered by real-time PCR. We did not observe any significant differences between the titered vectors **(Fig. 1b)**, suggesting that the above modifications did not affect the physical AAV packaging process. Next, titer-normalized vectors were transduced into HEK293T cells at MOI = 10,000. As shown in **Fig. 1c**, expression from vectors harboring Sp1 and NF-kB sites was significantly higher than that of the naïve EFS-NC counterpart.

Furthermore, the vectors carrying four repeats showed higher levels of NLuc expression compared to those with two repeats (**Fig. 1c).** The observed increase was close to 4-fold in the samples with 4 Sp1 or 4 NF-kB, and slightly higher increase in the sample bearing 2xSp1+2xNF-kB. Indeed, expression from the latter vector was only slightly lower than that shown by the vector carrying a complete copy of the EF1α promoter (**Fig. 1c).** This suggests that most of the enhancer activity provided by the distal (non-core) portions of the EF1α promoter is supplied by the binding of transcription factors Sp1 and NF-kB. The restoration of their binding sites within the expression plasmid had a major impact on the expression of the reporter transgene delivered by the viral vector.

### All-in-one AAV gene repression platform

Next, we set out to engineer a gene-silencing platform that fits within AAV size restrictions. The optimized cassette using 2 Sp1 and 2 NF-kB (see Fig. 1) was used throughout the remainder of this study. We utilized a screening strategy based on a dual reporter system, outlined in **Fig 2a**. A reporter vector expressing dGFP and NLuc genes was packaged into LV particles. The expression vector cassette also carried a puromycin resistance marker for selection. We then used HEK293T cells to create a stable reporter cell line. The cells were transduced at MOI = 0.2 to ensure integration at the rate of 1 copy per cell. The dGFP and NLuc proteins were expressed from the CMV promoter, which was targeted with gRNAs (**Fig. 2a & 2f**). Two gRNAs sequences were selected to target the reporter construct (**Fig. 2f**; gRNA1 and gRNA2). Importantly, both dGFP and NLuc are characterized by short protein half-lives, making them ideal for the evaluation of gene expression changes. In this study we focused on the catalytically inactive mutants of two small Cas9 proteins, derived from *Campylobacter jejuni* (dCjCas9) and *Staphylococcus aureus* (dSaCas9) (**Fig. 2a & 2f**).

**Fig. 2.**
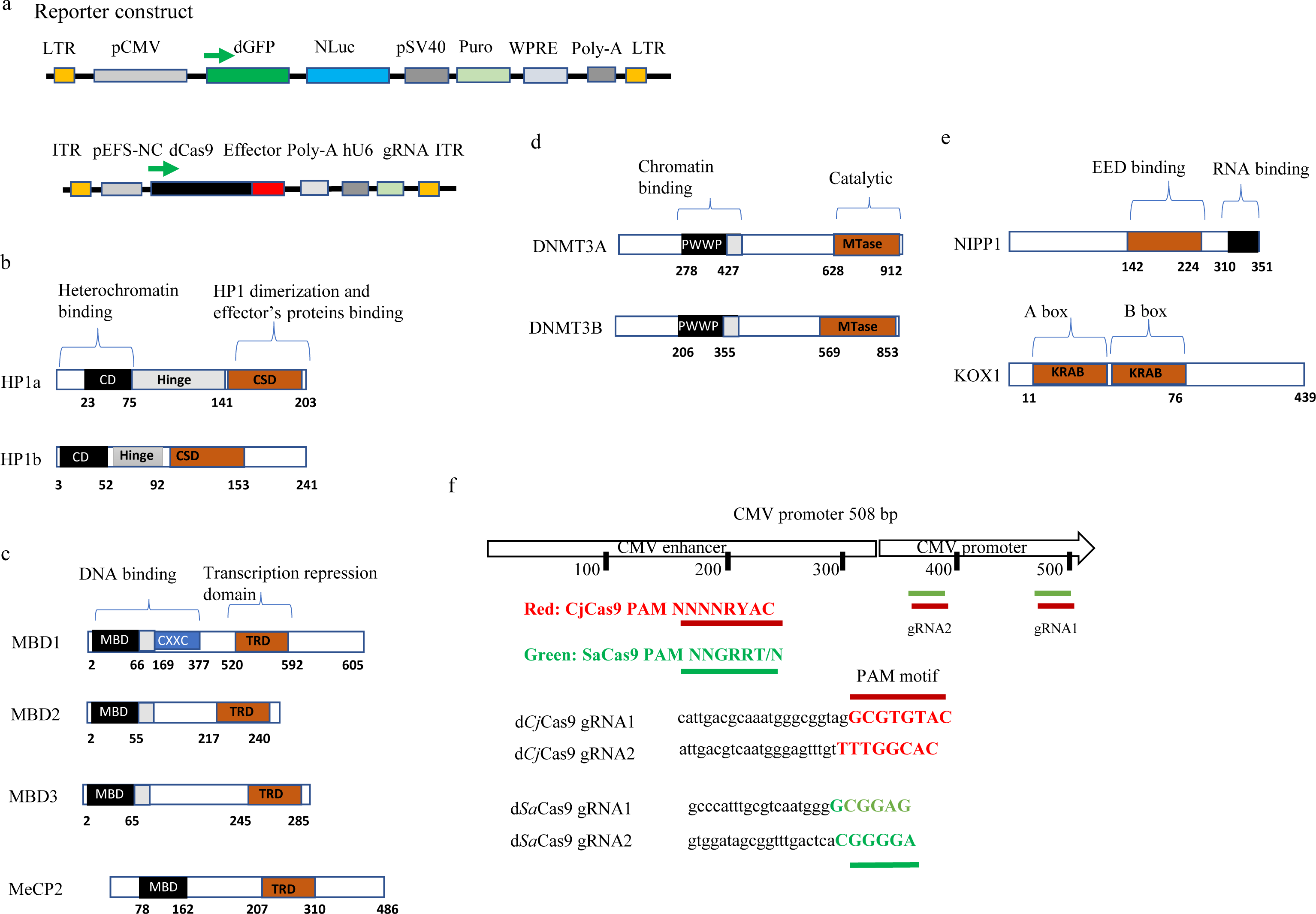

These proteins were then fused with a panel of repressors for screening: First, dCjCas9 and dSaCas9 were fused with heterochromatin proteins 1a and 1b (HP1a and HP1b, respectively) ([43] and **Fig. 2b)**. Here, we decided to use the repressive chromoshadow domain (CSD) of HP1 both with and without the hinge region. The chromoshadow domain is responsible for the multimerization of HP1 and the formation/maintenance of compact heterochromatin structures, while the hinge region is responsible for additional transcriptional regulation via interaction with histone H3 and other repressive effectors ([37], [44], and **Fig. 2b**). Secondly, we investigated TRDs from the methyl-CpG-binding domain (MBD) protein family **(Fig. 2c)**. This family contains Methyl-CpG-binding domain proteins 1, 2, and 3 (MBD1-3 respectively), and methyl CpG binding protein 2 (MeCP2). Note that Methyl-CpG-binding domain protein 2 (MDB2) and methyl CpG binding protein 2 (MeCP2) are different proteins **(Fig. 2c)**. Their respective TRDs are responsible for the repression mediated by these proteins [45]. As mentioned above, we previously developed a lentivirus carrying a dCas9-DNMT3A transgene for epigenome editing of the Parkinson’s risk-factor gene *SNCA* [19], [20], [46]. As such, a third group of repressors was also used in this study, based on DNA methyltransferases 3A and 3B (DNMT3A and DNMT3B) (**Fig. 2d)**. Lastly, we included a few miscellaneous repression domains: the TRD from Nuclear inhibitor of protein phosphatase 1 (NIPP1), a representative Krüppel associated box (KRAB) domain from Zinc finger protein 10 (KOX1), and a novel KRAB-MeCP2(TRD) combination (**Fig. 2e**).

The corresponding plasmids were packaged into AAV2.9 viral particles and concentrated as described in Materials and Methods, and the titers of the produced vectors were measured by real-time PCR. All vectors, except those carrying DNMT3A or DNMT3B, consistently achieved high physical titers **(Fig. 3a, b)**. This was not a surprise, as the dCjCas9-DNMT3A/B (∼5.1kb) and dSaCas9-DNMT3A/B (∼5.4kb) constructs are significantly oversized even for the optimized AAV backbone **(Fig. 3a, b)**. Notably, the physical titers obtained from the rest of the vectors produced here were similar to those obtained from a naïve vector carrying no CRISPR/Cas components ([23] and **Fig. 3a, b)**.

**Fig. 3.**
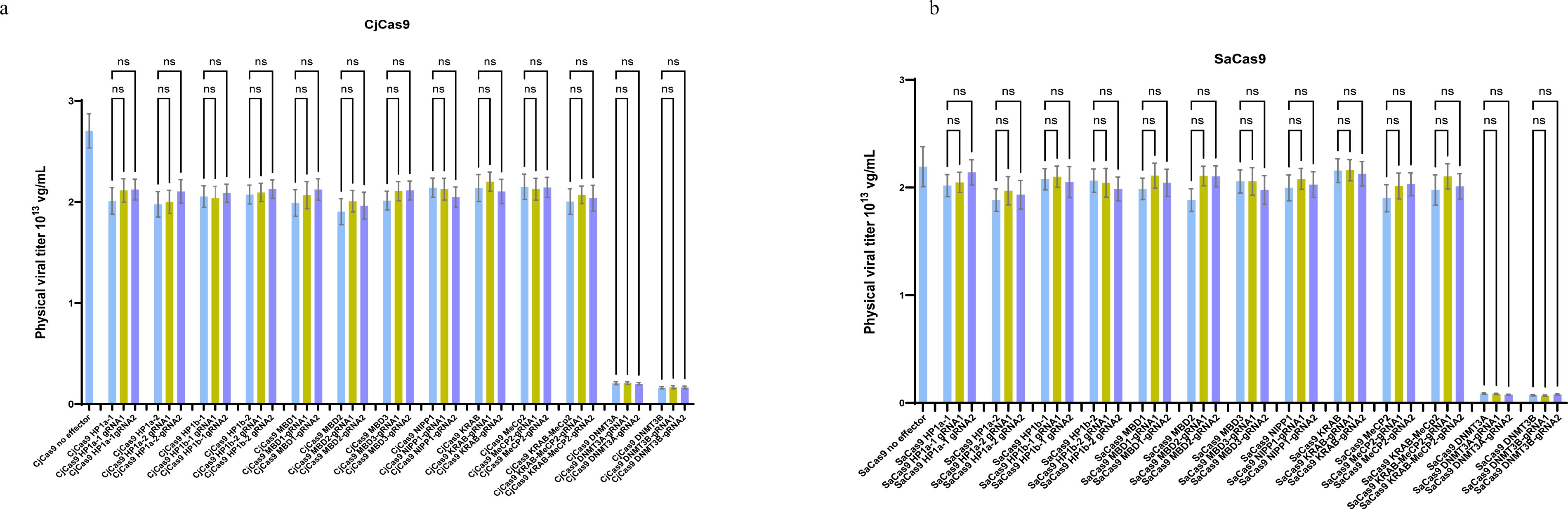
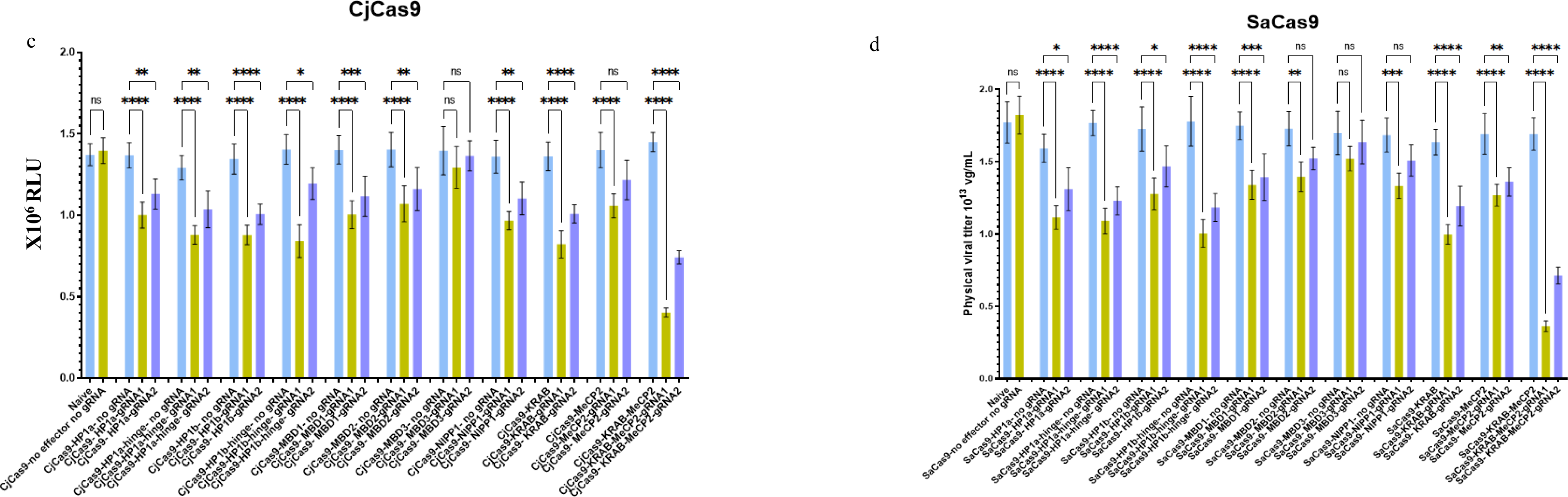
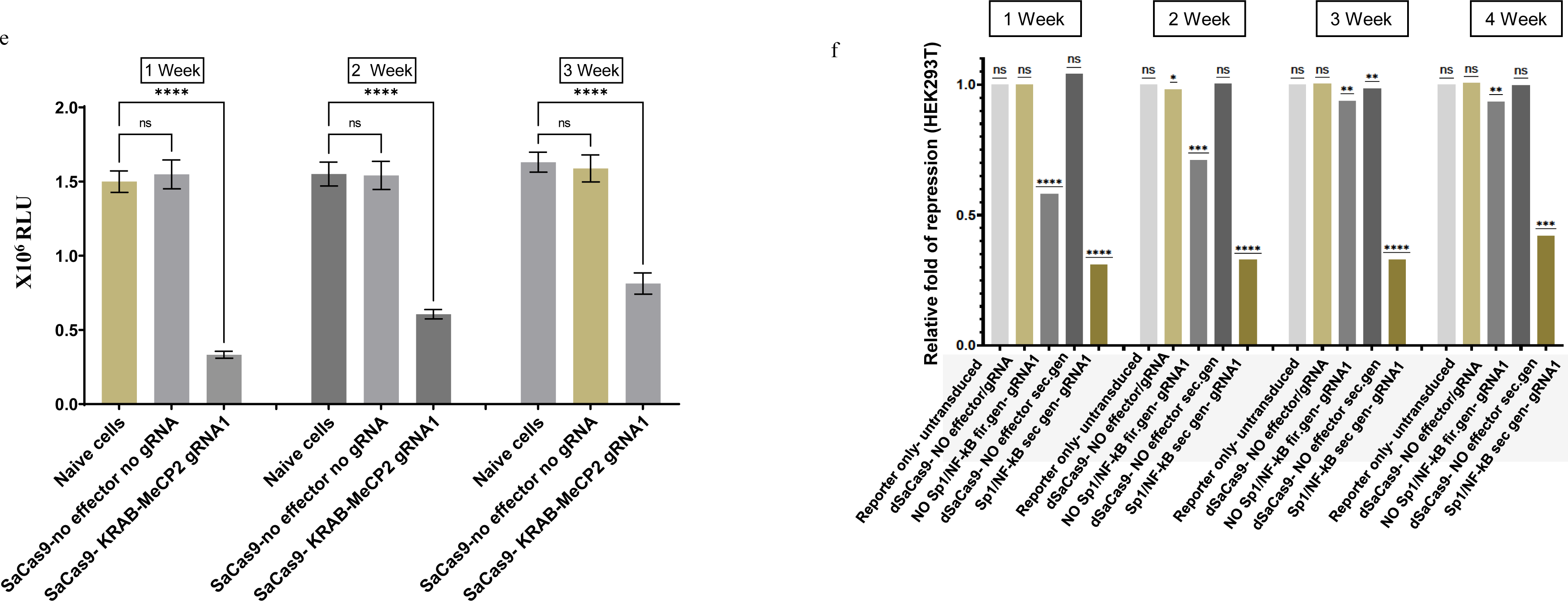
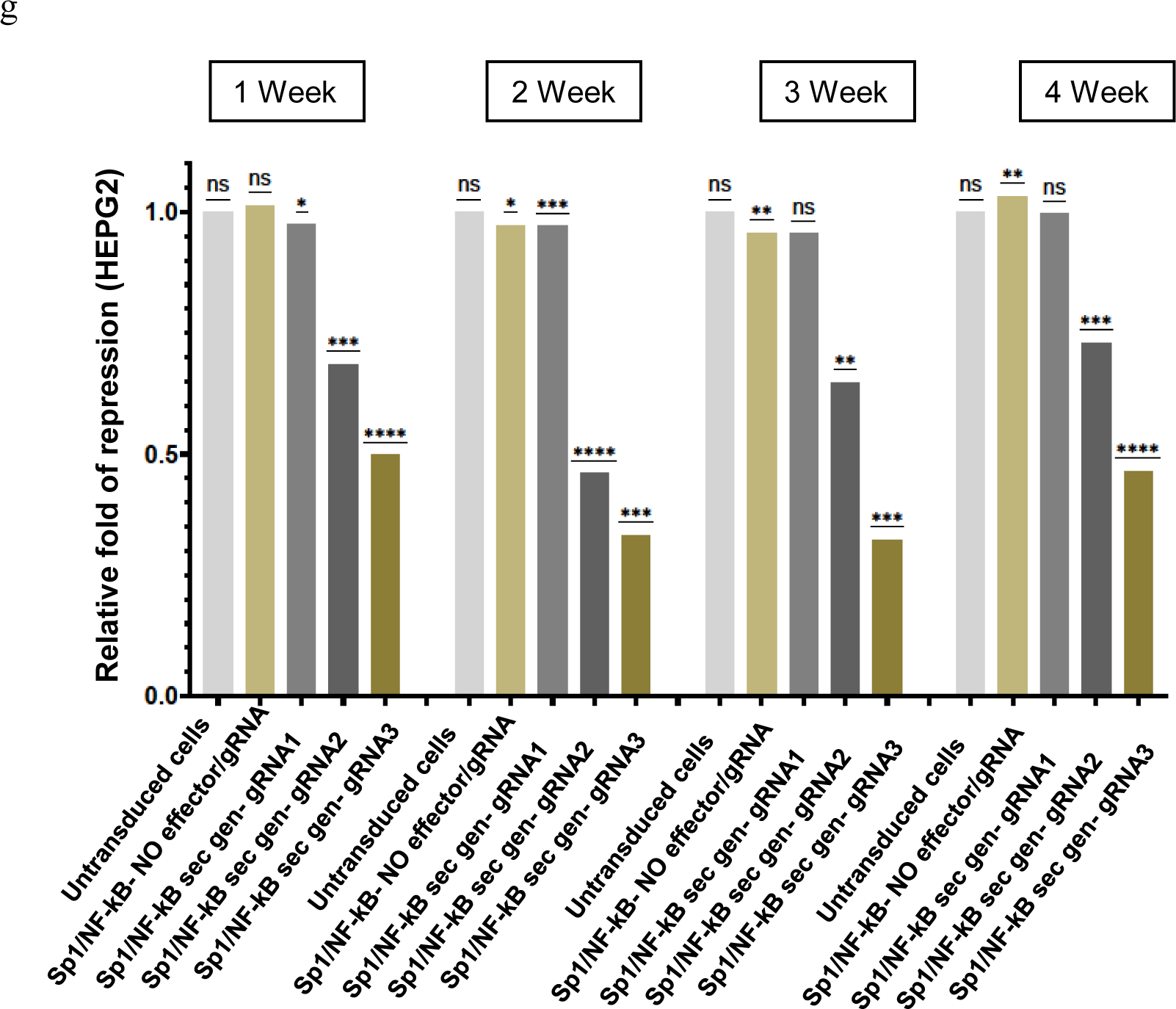

Next, the vectors were titer-normalized and transduced into the HEK293T reporter cell line (see Fig. 2a) to evaluate their repressive effects. The cells were harvested at day 4 post transduction, total protein was normalized via BCA assay, and a luciferase assay was performed. Importantly, no reduction in NLuc expression was detected in any of the samples transduced with vectors not expressing a CMV-targeting gRNA (across all effectors, plus a no-gRNA/no-effector double control), compared to naive control cells **(Fig. 3c, d)**. However, most samples transduced with vectors expressing dCj/SaCas-effector + [CMV gRNA] showed reduced NLuc expression (**Fig. 3c, d)**. Zooming in on the repression capacity of the selected effectors: HP1a and HP1b constructs containing the hinge in addition to the CSD performed slightly better than their CSD-only counterparts (**Fig. 3c, d)**. MBD1-, MBD2-, and NIPP1-containg constructs showed moderate levels of repression, but generally less than the HP1 family (**Fig. 3c, d)**.

Further, both Cas9 fusions with the MBD3 repressor were totally inactive. Finally, while the dCj/SaCas9-KRAB and dCj/SaCas9-MeCP2(TRD) constructs each individually resulted in significant levels of repression, their combination, dCj/SaCas9-KRAB-MeCP2(TRD), clearly demonstrated the most substantial silencing observed in the entire screen by a substantial margin (**Fig. 3c &d**). In fact, dCjCas9-KRAB-MeCP2(TRD)/[CMV gRNA1] reduced the level of NLuc expression by about 70%, and dSaCas9-KRAB-MeCP2(TRD)/[CMV gRNA1] lowered it by about 80%. The KRAB-MeCP2 vectors expressing CMV gRNA2 were less potent than those with gRNA1, but still effective: dCjCas9-KRAB-MeCP2(TRD) [CMV gRNA2] suppressed NLuc expression by about 50%, and dSaCas9-KRAB-MeCP2(TRD)/[CMV gRNA2] reduced it by about 60%. Interestingly, gRNA1 consistently outperformed gRNA2, showing greater reductions in luciferase signal across all effectors, not just KRAB-MeCP2(TRD) (**Fig. 3c, d**).

These results are consistent with earlier findings demonstrating that higher levels of repression are often achieved when gRNAs are designed to target a locus of interest in the vicinity of the transcription start site (TSS) [15], [24]. Indeed, gRNA1 targets a sequence which is proximal to the TSS, whereas gRNA2 targets an upstream part of the promoter (**Fig. 2f**).

### In vitro proof-of-concept

Whether epigenetic perturbations caused by episomally-expressed transgenes can be inherited and persist across generations of quickly dividing cells is not fully understood [24], [47]. Here, we aimed to analyze the dynamics of silencing and possible reactivation after an extra-chromosomally expressed AAV-CRISPR/Cas repression system is diluted out following cell divisions. To that end, we transduced HEK293T-derived reporter cells with AAV-gRNA1/SaCas9-KRAB-MeCP2(TRD) and extensively passaged them in the culture over the course of three weeks. The samples were collected at day 7, 14 and 21 post transduction, followed by BCA normalization and a Luciferase assay, as above **(Fig. 3e)**. As shown in **Fig. 3e**, transduction of the cells with the AAV-dSaCas9-KRAB-MeCP2(TRD) vector rapidly gave rise to gene silencing, which stably persisted through cell divisions. Consistent with the data reported in **Fig. 3c** and **d**, the level of NLuc repression was found to be ∼80% at day 7; and ∼60 and ∼50% after two and three weeks, respectively. These data suggest that the CRISPR-induced repression was readily transmitted across multiple cell divisions, resulting in heritable gene silencing. Next, we sought to complement this experiment by utilizing dGFP as a reporter readout. To that end, the HEK293T cells were transduced with AAV-gRNA1/SaCas9-KRAB-MeCP2(TRD) and extensively passaged them in the culture over the course of four weeks. Here we also used the previous generation of the AAV backbone, with no Sp1 and no NF-kB elements (**Fig. 3f**). The samples were collected at day 7, 14, 21 and 28 post transduction, followed by BCA normalization and a Western Blot assay. As shown in **Fig. 3f**, transduction of the cells with both the first gen vector (no Sp1/NF-kB) and the second gen vector (2Sp1-2NF-kB) rapidly gave rise to gene silencing at repression levels of 60% and 80%, respectively. However, only the vector harboring Sp1/NF-kB elements had a stable effect across cell divisions. Consistent with the data reported in **Fig. 3c-e**, the level of dGFP repression was found to be ∼70% at day 21 and ∼50% at day 28. Notably, the no-Sp1/NF-kB virus failed to generate significant repression at those time points.

These data suggest that the AAV-Sp1/NF-kB-gRNA1/SaCas9-KRAB-MeCP2(TRD) vector is capable of faithfully propagating transcriptional repression in HEK293T cells across multiple cell divisions, resulting in heritable gene silencing (**Fig. 3f**). To demonstrate the potential of the novel vector to modulate expression of clinically relevant targets, we used the developed an all-in-one AAV system to suppress transcription of the Pcsk9 gene in human liver hepatocellular carcinoma HEPG2 cells. Pcsk9 encodes a liver enzyme that regulates low-density lipoprotein (LDL) receptor degradation. Remarkably, loss-of-function mutations in Pcsk9 are associated with low serum cholesterol levels and reduced risk of cardiovascular disease with no noticeable side effects [48], [49]. Thakore and colleagues recently utilized a dual-vector AAV system to deliver dSaCas9-KRAB and a single gRNA for targeted repression of an endogenous Pcsk9 gene in mouse hepatocyte cell line and in vivo. Notably, the study reported that the Pcsk9 serum levels were reduced >90% over 4 weeks after treatment, but this magnitude of silencing was not sustained [29]. To test whether the transient AAV-Sp1/NF-kB-/SaCas9-KRAB-MeCP2(TRD) vector can permanently propagate the transcriptional silencing of the Pcsk9 gene, HEPG2 cells were transduced with vectors carrying three different gRNAs targeting the endogenous Psck9 gene (gRNA1-3), as well as a no-gRNA control vector, and extensively passaged them in the culture over the course of four weeks (**Fig. 3g**). Consistent with the results reported in **Fig. 3f**, the gRNA3-expressing vector demonstrated the characteristic stable silencing pattern that has been stably propagated through multiple cell divisions.

Using the gRNA3 expressing vector, Pcsk9 repression levels of ∼70% and ∼55% were measured at the 3- and 4-week time points, respectively. These effects were gRNA sequence-specific; a significantly lower level of repression was observed with the vector harboring gRNA2, and no noticeable repression was detected with gRNA1 (**Fig. 3g**). These results demonstrated that the novel system could facilitate long-term and stable gene silencing following a single application. Next, we assessed the integration capacity of the vectors to rule out the possibility that overexpressed CRISPR/dCas9 may alter the rate of integration. We transduced 293T cells with the first gen (no Sp1/NF-kB) and second gen (+Sp1/NF-kB) KRAB-MeCp2(TRD) vectors, as well as with a naïve AAV control vector. Then, as described above, we passaged the cells for 4 weeks to dilute out non-integrated viral genomes and finally isolated viral DNA from for analysis with real-time PCR. The integration rates were determined as a ratio between copy numbers at week 1-4 and at 24 hr post transduction (**Supplementary Fig. 1**). Consistent with our previous data, CRISPR/Cas9 components do not significantly alter the integration capacity of AAV vectors [22], [38]. Importantly, the data reported in **Supplementary Fig. 1** suggest that the episomal status of all-in-one CRISPR/dCas9 vectors is not compromised by the editing components, and that the stable mode of repression reported in Fig. 3 arises from transient expression of the editing machinery.

This suggests that when the rules governing the maintenance of different chromatin modifications are better understood, epigenome editing could be used as a potent ‘hit and run’ strategy for permanently modulating genetic loci, enabling long periods of drug-free state after the initial treatment. To test this hypothesis, a chromatin immunoprecipitation (ChIP) assay was used to determine whether the state of histone H3, one of the major four histones comprising the nucleosomal core, is associated with the repressive state established and maintained throughout cell divisions. Histone modifications such as acetylation of H3 and H4 and di- and trimethylation of H3 lysine 4 (H3-K4) have been associated with open chromatin and gene activation, while closed chromatin and gene repression is associated with trimethylation of H3 lysine 9 (H3-K9) [39]. To study the mechanism of the silencing mediated by the vector, a ChIP assay was used to determine which of the above histone modifications were associated with the CMV promoter of the reporter proviral DNA in HEK293T cells (**Fig.2a**) at day 7, 14, 21 and 28 pt. As shown in **Supplementary Fig. 2a**, the histone modification profile associated with the CMV promoter in the HEK293T-dGFP-NLuc cells transduced with AAV-Sp1/NF-kB-gRNA1/SaCas9-KRAB-MeCP2(TRD) vector was typical of transcriptionally repressed genes, showing low levels of H3 and H4 acetylation, H3-K4 dimethylation, and an enrichment of trimethylated H3-K9. Importantly, this repressive pattern was faithfully maintained throughout the experimental period (**Supplementary Fig. 2a-c**). Those results clearly demonstrate that the novel AAV system is enabling faithful propagation of the repressive state through multiple cell divisions. In contrast, the vector system that harbors no Sp1/NF-kB elements failed to maintain this repressive pattern at later time points, consistent with the transient mode of silencing reported in **Fig. 3c-f**. To evaluate whether similar permanent repressive chromatin marks could be found on endogenous Pcsk9 target gene, we used a ChIP assay on the HEPG2 cells described above, which had been given a single treatment with AAV-Sp1/NF-kB/SaCas9-KRAB-MeCP2(TRD) + Pcsk9 gRNA1-3. Consistent with the data reported in **Supplementary Fig. 2a-c**, we found the promoter of the Pcsk9 gene silenced via AAV-Sp1/NF-kB/SaCas9-KRAB-MeCP2(TRD) encased in heterochromatin, characterized by low levels of H3 and H4 acetylation and H3-K4 dimethylation, and enrichment of trimethylated H3-K9 (**Supplementary Fig. 3a-c**). Importantly, this repressive pattern was faithfully maintained throughout the experimental period of 4 weeks in HEPG2 cells as well. Consistent with the expression patterns demonstrated in **Fig. 3g**, the repressive marks were observed strongly with gRNA3 and to a lesser extent with gRNA2. Lastly, we did not observe any noticeable repressive modifications following transduction with gRNA1 or control (no gRNA) vectors, again highly correlated with their repressive capacities (**Fig.3g**).

### In vivo proof-of-concept

Studies using *in vivo* models to evaluate the efficiency, specificity, and safety of AAV delivered CRISPR/dCas9 platforms serve as the “gold standard” in the preclinical stages of drug development. To the best of our knowledge, no study thus far has reported successful epigenetic editing in animals using an all-in-one AAV platform. Here we aimed to validate our lead epigenome editing platform *in vivo*, in the mouse hippocampus. The hippocampus is the primary affected brain region in late onset Alzheimer’s disease (LOAD) and related diseases (reviewed in [50]). Therefore, we chose this region to assess the potential utility of our platform as a potential system for LOAD therapy.

First, to evaluate the efficacy of the platform, we stereotaxically co-injected the AAV/dSaCas9-KRAB-MeCP2(TRD)/gRNA1 repressor with the LV/dGFP-nLuc reporter (hereafter, LV/GFP) into the left dorsal hippocampus (DH). In parallel, we co-injected the no repressor/no gRNA control construct (AAV/dSaCas9) with the LV-GFP reporter into the right DH, such that each mouse provided an internal control. Brains were harvested at 2 timepoints post-injection; 14 days or 42 days. GFP protein and RNA expression was compared between the left and right DH for each mouse. This experiment was performed using C57BL/6 mice of 2 age groups; were young adults (16 weeks) and middle-aged (32 weeks old).

There was no significant difference in GFP expression between males and females, thus, the data was combined for subsequent analysis. In young adult mice, injection of the AAV/dSaCas9-KRAB-MeCP2(TRD)/gRNA1 vector significantly repressed GFP expression in the left DH compared to the control right DH at both 14 days and 42 days after injection in the young adult group (14d: 45.18% ± 7 reduction t(8)=4.194 p=0.003, paired *t*-test. 42d: 48.06% ± 11 reduction, t(10)=3.944 p=0.028, paired t-test. **Fig. 4a, b**)., Consistently, GFP mRNA expression levels were also significantly repressed by the AAV/dSaCas9-KRAB-MeCP2(TRD)/gRNA1 vector after both 14 days and 42 days in the young adult group (14d: t(4)=3.403 p=0.027, paired t-test. 42d: t(5)=4.505 p=0.0064, paired t-test. **Fig. 4c**). There was no significant difference in the efficacy of the repressor between the two incubation durations. Injection of the AAV/dSaCas9-KRAB-MeCP2(TRD)/gRNA1 vector also significantly repressed GFP expression in the middle age group (59.02% ± 6 reduction, t(9)=4.116 p=0.026, paired t-test–; **Fig. 4a, b**). There was no significant difference in the efficacy of the repressor between the two age groups. Collectively, the AAV/dSaCas9-MeCP2(TRD)/gRNA1 repressor effectively reduced GFP reporter expression by ∼45-60% in mouse DH.

**Fig. 4.**
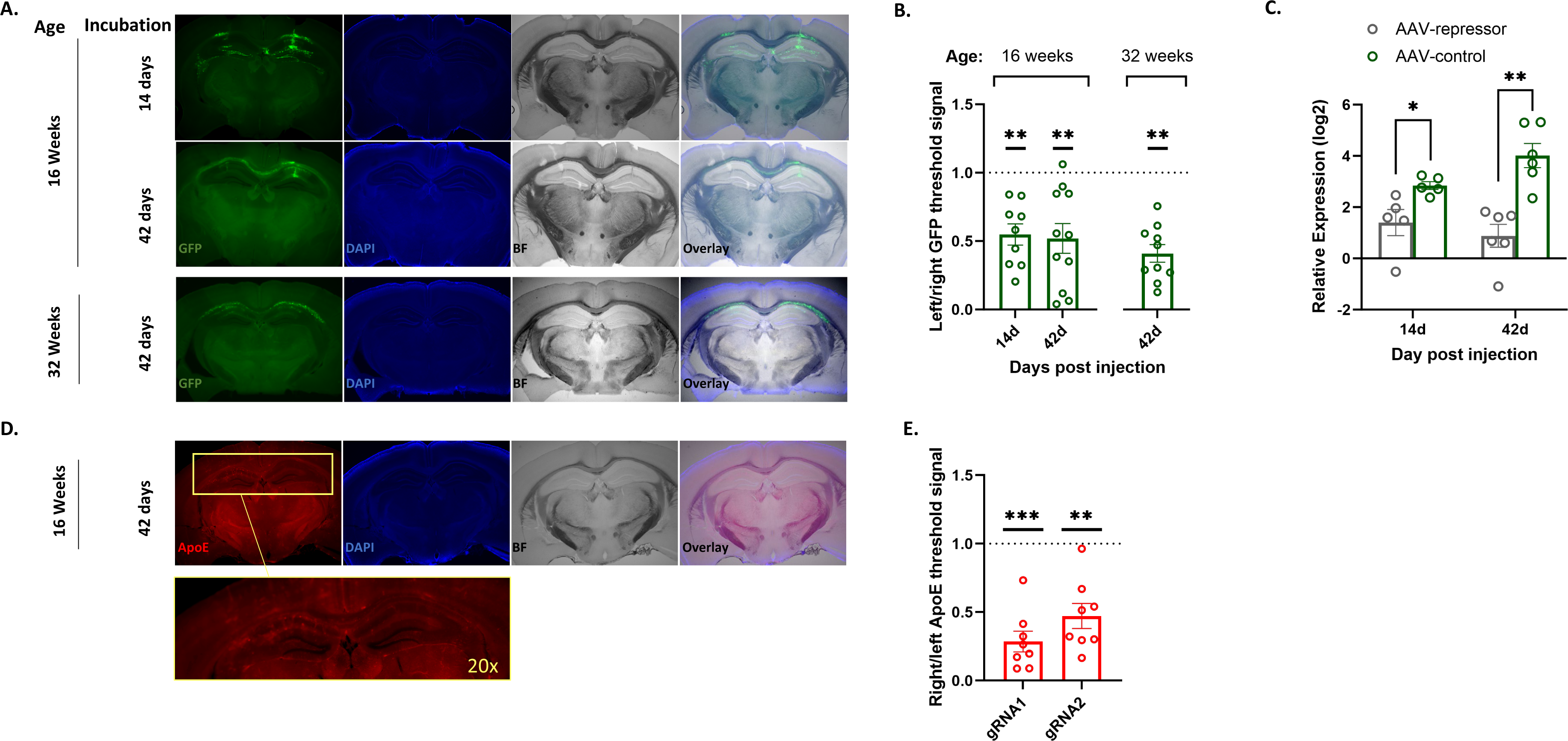

*APOE* is the strongest and most reproducible genetic risk factor for LOAD, and an emerging therapeutic target for the disease (reviewed in [51]). Thus, we next investigated the *in vivo* efficacy of our all-in-one AAV epigenome-editing platform using *APOE* as a therapeutically relevant target for LOAD. Similarly to the reporter experiment described above, we chose the DH as - a brain region affected in early stage of LOAD. However, in this experiment, we targeted the expression of the mouse endogenous *Apoe* in middle-aged C57BL/6 mice Two all-in-one AAV repressor vectors were used, which carried dSaCas9-KRAB-MeCP2 along with two different gRNAs targeting the mouse *Apoe* promoter and the efficacies in reducing its expression were tested and compared. A total of sixteen mice (8 males, 8 females) received bilateral stereotaxic injection with the AAV/dSaCas9-KRAB-MeCP2(TRD)/[*Apoe*-pro gRNA] repressor vector carrying either gRNA1 (n=8) or gRNA2 (n=8) into the right DH, and the AAV/dSaCas9 control vector (no gRNA, no repressor) into the left DH. At six weeks post-injection, the expression of mouse ApoE protein was quantified by immunohistochemistry and the levels in right DH relative to the left DH were determined for each mouse. Again, there was no significant difference in *Apoe* expression between males and females, thus, the data was combined for subsequent analysis. The results showed that both AAV/dSaCas9-KRAB-MeCP2(TRD)/[*Apoe*-pro gRNA] vectors significantly repressed endogenous mouse ApoE expression in the right DH compared to the left DH (gRNA1 – two-tailed Mann-Whitney U = 0, n = 8, P = 0.0002; gRNA2 – two-tailed Mann-Whitney U = 3, n = 8, P = 0.0011; **Fig. 4d, e**). The relative expression levels of ApoE (right DH/left DH) with gRNA1 and gRNA2 were 28.8% ± 7.07 and 47.16% ± 8.5, respectively (**Fig. 4d, e**). However, there was no significant difference in the efficacy between the two gRNAs (Unpaired t-test, p=0.14). In conclusion, the endogenous expression of mouse ApoE was significantly and robustly reduced by ∼71% and ∼53% by our all-in-one repressor platform carrying gRNA1 and gRNA2, respectively. These results indicated that our AAV-delivered epigenomic therapy platform can effectively suppress ApoE expression, and thus bears translational potential towards the development of therapeutics approach to treat, prevent, and/or delay the progression of LOAD.

Finally, we conducted pilot safety evaluations including daily weights of the treated mice and monitoring welfare criterions. In the GFP reporter experiment, while both age and sex had a significant influence on mouse weight (young adult – Male = 26.09 ± 0.06, Female = 22.37 ± 0.1, F(1,10) = 28.97, p = 0.0003; middle age – Male = 32.17 ± 0.09, Female = 25.43 ± 0.05, F(1,10) = 37.48, p = 0.0001; two-way ANOVAs), all mice weights were stably maintained during the 42 days between viral vector injection and sacrifice(**Fig. S3a**). Similarly, in the endogenous *Apoe* experiments we observed no changes in the weights of the mice during the 42 days following vector injection (**Fig. S3b**). In addition, the mice displayed normal grooming and eating/drinking behaviors. Overall, these analyses did not reveal any safety issues after the stereotaxic injection of our all-in-one AAV platform for gene repression mediated by epigenome editing.

## Discussion

In this study we applied a screening strategy using a lentiviral dual reporter system to identify dSaCas9-KRAB-MeCP2(TRD) as being the most potent and sustainable Cas9-effector combination for epigenome-mediated repression. Our results support a recent study by Yeo and colleagues, who engineered a highly effective dCas9-KRAB-MeCP2 transcriptional repressor [32]. Nevertheless, their final vector design contains both the complete MeCP2 ORF and the bulky SpCas9 and would be vastly oversized for packaging into AAV particles. More recently, a similar expression cassette was re-cloned into a lentiviral backbone, yielding lenti_dCas9-KRAB-MeCP2 (Andrea Califano, unpublished data; Addgene plasmid #122205). This vector also carries the complete MeCP2 cDNA and dSpCas9. Here, we report the development of a miniature repressor system using only the MeCP2 transcriptional repression domain (TRD) [52], [53], [54], [55], [45]. The TRD identified in the above studies was sufficient to mediate robust, sustained gene silencing by providing a binding platform for HDAC1/2 and other transcriptional repressors. The TRD that was used in this study comprised amino acid residues 207-310 of MeCP2. The coding sequence of the dSaCas9-KRAB-MeCP2(TRD) transgene was only 3.6 kb long, allowing us to include promoter, transgene, polyA signal, U6-gRNA cassette, and ITRs, and remain narrowly within the AAV packaging capacity. With this vector we achieved physical titers of slightly over 10^13^vg/mL, which is a sufficient titer for most therapeutic applications in humans. Most importantly, the vector demonstrated a high level of gene repression both *in vitro* and *in vivo* when transduced into cultured cells or mouse hippocampus, respectively.

The efficiency and specificity of the epigenome-editing approach has shown great promise for a wide range of gene therapy applications. Initial successes have been reported in many studies and have inspired efforts to improve and optimize CRISPR/dCas systems for targeting and manipulating DNA at the epigenetic level (reviewed in [24]). These systems bear significant advantages over conventional CRISPR/Cas editing approaches utilizing active Cas enzymes, including (*i*) higher specificity and on-target editing efficiency, (*ii*) lower off-target effects, and (*iii*) an inability to break – or even nick – DNA at the site of interest, resulting in lower toxicity. Similarly, rates of undesired genomic rearrangements such as deletions, duplications, inversions, and translocations are significantly lower when using dead Cas9-based editing systems [15], [31]. Combining CRISPR/dCas9 targeting components with potent epigenetic repressors in a single AAV delivery vector will greatly facilitate studies of gene regulation and the development of new approaches to address gene dysregulation in various disease states.

Adeno-associated vectors (AAVs) represent the “gold-standard” delivery platform for a range of gene therapy applications [23]. Their high efficiency, capacity to robustly transduce both dividing and non-dividing cells, and the ability to persist in non-integrating/transient forms have elevated AAVs to be the platform of choice for applications in both basic and translational research. However, there is a challenge: AAVs can only carry relatively small transgenes, whose size does not exceed approximately 4.7kb [21]. Several approaches have been developed to circumvent this bottleneck. First, an *in situ* split intein-based approach has been used in which the coding sequence of Cas9 is split in half, delivered via two AAVs, and the full protein is reassembled after transduction [56], [57]. The second approach utilizes a far simpler type of dual-vector system, in which the two components of the repression machinery are delivered from separate AAVs (dCas/effector from one AAV, gRNA cassette from the other) [56], [58], [29]. Although successes have been reported, these approaches usually demonstrate only moderate efficacy, and require higher viral doses to achieve the desired effects. Furthermore, manufacturing of dual-AAV systems is time- and cost-intensive. These shortcomings served as the impetus for our development of an AAV platform capable of efficiently transferring fully functional CRISPR/dCas repressor complexes to a cell or tissue of interest in a single AAV particle. Our all-in-one vector accomplishes exactly that, defeating AAV packing limitations for the first time. As such, our platform is attractive and advantageous for both basic and translational sciences as well as clinical applications.

It is not currently clear whether transcriptional changes driven by epigenome editing can be inherited in a stable and permanent manner. On the same note, it is not well understood whether epigenetic modifications introduced by epigenome editors will be ‘remembered’ and propagated by human cells without constitutive and integrated expression of the epigenetic modifier. In fact, it has been demonstrated that programmable epigenetic tools expressed transiently are tunable and reversible [59]. In contrast, other studies have shown that it is feasible to install a stable transcriptional program that is inherited across cell divisions, without integrated expression of the epigenetic modulators [60], [61]. In particular, Amabile et al. demonstrated heritable silencing by recruitment of DNA methyltransferases and KRAB proteins to the target genes. However, this and other approaches aiming to permanently shut down expression of a target of interest are based on a design utilizing either two or three fusion proteins for each gene, which is experimentally cumbersome, especially for multiplexed gene targeting. Furthermore, the multiplexed repressors used in the above studies consistently cannot be delivered by AAV, being far over the size restrictions. Similarly, a TALE-based fusion of KRAB and the DNMT3A and DNMT3L domains resulted in low efficacy but long-term gene repression [62]. A recent study conducted by Thakore and colleagues also demonstrated a transient gene silencing effect [29]. This study used a dual AAV vector system (separately carrying dSaCas9-KRAB and the targeting gRNA) for gene silencing of Pcsk9, a regulator of cholesterol levels, in the liver of adult mice. Systemic administration of the dual-vector AAV8 system resulted in significant reductions of serum Pcsk9 and cholesterol levels, however this reduction was not sustained [29]. In contrast with those findings, here we demonstrate that significant repression (∼50%) of a reporter gene persists for at least four weeks post-transduction in fast-dividing HEK293T cells. By this time point, the cells had divided more than 20,000 times and carried no detectable viral genomes, suggesting that transient expression of dSaCas9-KRAB-MeCP2(TRD) can cause robust, sustained gene silencing that is ‘remembered’ and inherited by cells through multiple cell divisions. As such, the mitotically stable ‘hit-and-run’ epigenome editing approach developed in this study is extremely appealing for correcting dysfunctions in dividing or non-dividing cells, in both *ex vivo* and *in vivo* settings. Even more broadly, the viral system developed here can be adjusted to target a cell or tissue of interest by selecting the desired viral serotypes or perhaps outfitting the epigenome modifier with a cell-or tissue-specific promoter (if short enough). And of course, the CRISPR/Cas system can target any sequence of interest by simply switching the gRNA protospacer sequence. Finally, additional regulation may be achievable by our developed platform via repressor and/or gRNAs multiplexing, adding a substantial advantage.

As mentioned above, the developed platform is highly efficient for modulating gene expression in both *in vitro* and *in vivo* models. To evaluate the efficiency of the developed system, we utilized the HEK293T-reporter cell line and demonstrated that a gRNA targeting the core (TSS-proximal) part of the CMV promoter resulted in an approximately 80% reduction in reporter expression. Furthermore, we validated the developed vector on two therapeutically relevant targets. To that end, we demonstrated that AAV9 delivery of gRNA/dSaCas9-KRAB-MeCP2 to human liver hepatocarcinoma HEPG2 cells resulted in a potent, sustained, and durable silencing of the cholesterol regulator Pcsk9 in the liver of adult mice [48], [49]. Permanent gene regulation with dCas9/CRISPR*i* is typically achieved with stable expression of the gRNA and dCas9-effector proteins; this is usually achieved via integrase-competent lentiviral vectors (ICLVs). In contrast, here we report that an all-in-one, transient AAV gene delivery system provided long-term episomal expression with minimal genomic integration in fast-paced dividing cells such as HEPG2 and HEK293T. In fact, the developed system has proven capable of reliably propagating the transcriptional silencing in both rapidly dividing cells in culture and *in vivo* in non-dividing, post-mitotic cells in the CNS (see below in the discussion), throughout an extended experimental period in both cases. Consistent with this observation, we found that post-translational histone modifications associated with open chromatin and gene activation, such as acetylation of H3 and H4 and di- and trimethylation of H3 lysine 4 (H3-K4) were significantly depleted in the cells transduced with the repressor-expressing vector, while the characteristic marker of closed chromatin and gene repression H3 lysine 9 (H3-K9) trimethylation was substantially enriched. Importantly, we demonstrate that the episomal status of AAV-CRISPR*i* vectors remains uncompromised, and that the sustained gene silencing reported in **Fig. 3** arises from transient expression of the epigenetic effector.

In this study, we showed that the novel dSaCas9-KRAB-MeCP2 platform can be efficiently packaged into an all-in-one AAV particle delivered efficiently *in vivo* into the mice brain. Using the above CMV-driven dual-reporter system (expressed dGFP and Nano-Luciferase genes), we demonstrated ∼50-60% reduction in the expression of an exogenous dGFP reporter. Furthermore, an efficient and sustainable gene repression was observed between the time points of two- and six-weeks post stereotaxic injection of AAV9/dSaCas9-KRAB-MeCP2(TRD) into the mouse hippocampus. Most importantly, the durable repression by targeting the CMV promoter was observed with the virus that carried the optimized backbone, harboring 2xSp1 and 2Xnf-kB transcriptional activation sites. It has to be mentioned, that the similar approach to improve CRISPR/Cas9 delivery has been recently reported by our group. In the previous work, Ortinski and colleagues showed that the addition of Sp1 and NF-kB binding site into an episomal, integrase-deficient lentiviral vector (IDLV) results in a dramatic increase in packaging efficiency and expression [22], [63]. Further, we most recently developed a similar platform using a lentiviral vector (LV) backbone for neuronal type-specific epigenome editing aiming to decrease expression of the Parkinson’s disease risk factor *SNCA*. These studies support the therapeutic potential of our repressive platforms for Parkinson’s disease and dementia with Lewy bodies (DLB) [64], and provide the foundation for preclinical studies in animal models toward investigational new drug (IND) status and clinical trials. However, the platform reported in this work is by far the most promising, being to the best of our knowledge the first system to deliver all the necessary components for epigenome editing in a single AAV.

Most of the available vector expression cassettes lack Sp1 binding sites, and neither RNA Pol III promoters (e.g., U6 and H1, typically used to express sgRNA), nor viral core promoters (e.g., EFS-NC, expressing the dCas9-repressor), harbor Sp1 or NF-kB binding sites [22], Here, we clearly demonstrate that the addition of Sp1 and NF-kB sites to a relatively weak core promoter like EFS-NC can substitute for the use of the more powerful but larger full-size counterpart, here the full EF1-alpha promoter. Based on our results, we speculate that insertion of transcription activation binding sites could be adopted as a universal approach to enhance production, transcription, and infectivity of other episomal viral systems used for delivering CRISPR/Cas9 cargoes, or potentially any large transgene which puts promoter space at a premium.

Adopting the AAV-KRAB-MeCP2(TRD) system for use *in vivo* would facilitate studies of gene regulation in higher organisms and the development of approaches to tackle aberrant gene regulation in various disease states [24]. For those therapeutic applications, the AAV delivery platform is particularly advantageous as it has been extensively manufactured to target a variety of tissue and organ types, including the CNS [23]. Indeed, the potential benefits of using AAV delivery methods paired with epigenome editing are enormous as they share several key properties including low immunogenicity, lack of oncogenicity and pathogenicity, efficient gene transfer, long-term gene-of-interest expression, and scalable manufacture for clinical applications. Within the past 5 years, the gene therapy field has seen a wave of drugs based on AAV delivery platform that have gained regulatory approval for a variety applications [23], [15], [24]. To demonstrate the therapeutic utility and applicability of the platform, we performed validation experiments in the context of CNS disorders and specifically dementias. The dorsal hippocampus (DH) was selected as it plays a major role in learning and memory and its atrophy is one of the most consistent features in several age related neurodegenerative diseases (NDDs), including Alzheimer’s Disease (AD), Huntington’s Disease (HD), Frontal-Temporal Dementia (FTD) and Amyotrophic Lateral Sclerosis (ALS) (reviewed in [50], [51], [65]). We provided the example of Apolipoprotein E (*APOE*) gene as it is the strongest and most reproduceable genetic risk factor for late-onset Alzheimer’s disease (LOAD risk and age-at-onset (AAO) [66–79], and thus holds promise as a potential therapeutic target for LOAD and related dementias (reviewed in [51]). Furthermore, accumulating evidence suggests that increased overall expression of *APOE* plays an important role in the etiology of LOAD (reviewed in [51, 80]). Therefore, we applied our platform to evaluate its potential for downregulating *APOE* gene expression. We demonstrated that levels of endogenous mouse ApoE expression were reduced up to 70% following stereotaxic injection of the AAV/dSaCas9-KRAB-MeCP2(TRD) platform in the mice DH. In addition, our data supports the safety of the platform *in vivo* upon administration into the mouse brain These outcomes warrant further preclinical investigations in Alzheimer’s disease models. Collectively these results suggest that our innovative platform could serve as a promising foundation for the development of a disease modifying therapy (DMT) to prevent, delay the onset of, and/or halt the progression of LOAD. Moreover, the ability to quickly and easily tailor our platform to target genes associated with other neurodegenerative diseases such as *SNCA* in PD, DLB and MSA, broadens the applications of the platform to a wide range of CNS disorders caused by the dysregulation of any number of genes.

## Materials and Methods

### Plasmid design and construction

The *Cj*Cas9 was derived from the plasmid pX551-CMV-*Cj*Cas9 (addgene, #107035; gift from Alex Hewitt’s lab). The plasmid was amplified using the following primers: 1097AgeI-For, 5’-agctctctggctaactac-3’ and 1097BamHI-Rev, 5’-cttttattgGatCcttagctggcctcc-3’. The plasmid pBK694 was previously created in the lab and harbors *Sp*Cas9 in an AAV backbone. The EFS-NC promoter was used to drive expression of *Sp*Cas9 in the above backbone. The plasmid was digested with AgeI-BamHI and cloned with the corresponding fragment containing *Cj*Cas9. The resulting plasmid was named pBK1119. We then created the pBK1120 plasmid by replacing the BsmI-PmlI fragment with its counterpart carrying the H559A mutation in the *Cj*Cas9 ORF. The corresponding fragment was synthesized using GenScript synthesis service. The second mutation, inactivated *Cj*Cas9 D8A was introduced into pBK1120 via digestion and replacement of an AgeI-PflMI fragment created using GenScript synthesis service. The resulting plasmid, pBK1124, carried catalytically inactive CjCas9 (dCas9). Then, we created a CjCas9-U6-promoter-gRNA scaffold using the corresponding fragment synthesized via GenScript synthesis service. Next, we introduced a linker sequence containing DraI-SphI-BlpI 41bps site in frame with the CjCas9 protein. The resulting plasmid, pBK1294a, has been used as a common intermediate for the following cloning of all effector peptides fused with d*Cj*Cas9. The repressor-effectors were cloned into SpeI-BmtI restriction sites of pBK1294. The sequences of the repressors can be found in supplementary fig 1. To create SaCas9-based constructs, we used pX603-AAV-CMV:NLS-dSaCas9(D10A,N580A)-NLS-3xHA-bGHpA that was obtained from addgene (plasmid #61594; gift from Fang Zhang’s laboratory). The amplified fragment of 3202bp harboring dCas9 CDS was cloned into pBK694 as described above. The primers contained AgeI-BamHI sites which were used for cloning, as above. The resulting plasmid was pBK1124. The oligo flanked by NdeI-NotI sites was annealed and cloned to create the pBK1129 plasmid. We then amplified the C-terminus of *Sa*Cas9 and replaced it with the mutated version that carried no stop codon. The following oligo was used 5’-ggatcctcaaataaaagatctttgttttcattagatctgtgtgttggttttttgtgtgcggccggtacc-3’ to remove the stop codon at the C-terminus of *Sa*Cas9. This plasmid was named pBK1198. Then, we introduced the adaptor sequence downstream from C-terminus of *Sa*Cas9. The sequence was 5’-GATCCggtggaggaagtggcgggtcagggtcgggtggcACTAGTataGCTAGCggaggtggttcgccaaagaagaaacggaaggtg G-3’. The resulting plasmid is pBK1294b. This plasmid has been used as the intermediate vector for cloning all of the d*Sa*Cas9-repressor fragments. The following gRNA oligos targeting the CMV promoter were selected: *Cj*Cas9-to-CMVp-1 5’-cattgacgcaaatgggcggtag-3’ *Cj*Cas9-to-CMVp-2 −5’-attgacgtcaatgggagtttgt-3’. *Sa*Cas9-to-CMVp-1 5’-gcccattgacgcaaatgggc-3’; *Sa*Cas9-to-CMVp-2 5’-gtggatagcggtttgactca-3’.(see, also in Fig. 2f). The following gRNA oligos targeting the ApoE promoter were selected: *Sa*Cas9-to-(Apoe)-1 (derived from pBK1861) 5’-gaggagggggcgggacagg-3’; *Sa*Cas9-to-(Apoe)-2 (derived from pBK1863) 5’-gtagctcttccctcccaaggt-3’. pBK533-pLV-EFS-NC-GFP-P2A-Nluc-WPRE was created previously [22]. The MluI-EcoRI fragment containing EFS-NC-*e*GFP-p2a-nLuc from pBK533 was re-cloned into an AAV backbone. The resulting plasmid was named pBK1083. The pBK1034-lentiviral vector plasmid harboring two Sp1 binding sites was used for the subcloning of 2xSp1-EFS-NC-*e*GFP-p2a-nLuc into the AAV backbone. The resultant plasmid, pBK1084, carries 2xSp1 5-ggatcc**GGGCGGGAC***GTTAACGG***GGGCGGAAC**-3’(marked in bold); separated by the linker sequence (marked in italics). The pBK1035-lentiviral vector plasmid harboring four Sp1 binding sites has been used for the subcloning of 4xSp1-EFS-NC-*e*GFP-p2a-nLuc into the AAV backbone. The resulting plasmid, pBK1085, carries 4xSp1 5’-ggatcc**GGGCGGGAC***GTTAACGG***GGGCGGAACGGGCGGGAC***GTTAACGG***GGGCGGAAC**-3’. The Sp1 sites are marked in bold and red; the linker sequences are marked in italics. The pBK1036-lentiviral vector plasmid harboring two NF-kB binding sites has been used for the subcloning of 2x NF-kB-EFS-NC-*e*GFP-p2a-nLuc into the AAV backbone. The resulting plasmid, pBK1086, carries 2x NF-kB. 5’-ggatcc**GGGGACTTTCC***GTTAACGC***GGGGACTTTCC**-3’. The NF-kB sites are marked in bold; the linker sequence is marked in italics. The pBK1037-lentiviral vector plasmid containing four NF-kB binding sites was used for the subcloning of 4x NF-kB-EFS-NC-*e*GFP-p2a-nLuc into the AAV backbone. The resulting plasmid, pBK1087, carries 4x NF-kB. 5’-ggatcc**GGGGACTTTCC***GTTAACGC***GGGGACTTTCCGGGGGACTTTCC***GTTAACGC***GGGGAC TTTCC**-3’. The NF-kB sites are marked in bold; the linker sequences are marked in italics. The lentiviral vector plasmid pBK1038 harboring 2xSP1 and 2xNF-kB binding sites was used for the subcloning of 2xSp1 and 2xNF-kB-EFS-NC-*e*GFP-p2a-nLuc into the AAV backbone. The resulting plasmid, pBK1088, carries 2xSp1 and 2xNF-kB. 5’-ggatcc**GGGCGGGAC***GTTAACGG***GGGCGGAAC**GTTAACGGGCGTA**GGGGACTTTCC***GTTAAC GC***GGGGACTTTCC**-3. The Sp1 and NF-kB sites are marked in bold; the linker sequences are marked in italics. We derived the EF1a promoter from pBK814-pLenti-EF1a-GFP-p2a-Nanoluc-WPRE. The promoter was cloned into pBK533 to create pLV-EF1a-GFP-P2A-Nluc-WPRE. The resulting vector pBK573 has been recorded. pBK573-lentiviral vector plasmid harbored EF1a-GFP-p2a-Nanoluc-WPRE has been used for the subcloning of EF1a promoter into the AAV backbone. The resulting vector pBK1184 has been recorded. To construct the reporter plasmid used in this study for the screening of the epigenetic repressors, pBK59, an empty LV vector was cloned with BamHI-SalI fragments carrying destabilized dGFP. dGFP was derived from the Lentiviral-TOP-dGFP-reporter (addgene plasmid #14715; gift from Dr. Tannishtha Reya’s laboratory). NLuc was subcloned from pBK533-pLV-EFS-NC-GFP-P2A-Nluc-WPRE as described above. The resulting plasmid, pBK1340, is schematically highlighted in Fig. 2a. The following gRNA were used to target Pcsk9 promoter in HEPG2 cells: gRNA1-gacgtctttgcaaacttaaaac; gRNA2-ccttccagcccagttaggattt; gRNA3-ccgaaacctgatcctccagtcc.

### AAV Vector Production

Plasmids were all packaged into AAV9. AAV vectors were generated using a triple transient transfection protocol in HEK293T (ATCC® CRL3216™) human embryonic kidney cells using polyethyleneimine (PEI). Briefly, for each virus, media for four 25 mm plates of HEK293T cells at 70–80% confluency was first changed into Dulbecco’s Modified Eagle Medium (DMEM) (Gibco #: 11965-092) without fetal bovine serum (FBS) (Hyclone #: SH30087.01). One hour after triple transfecting the cells with pHelper (12 ug/plate), pAAV *Rep-Cap* (10 ug/plate), and pAAV ITR-expression (6 ug/plate) plasmids using PEI and DMEM, 10% FBS was added to the media. to a final concentration of After 72 hours, cells were collected, and the cell pellets underwent 4 freeze thaw cycles in −80°C. They were sonicated before being purified using two rounds of ultracentrifugation on a cesium chloride (CsCl) gradient. After each centrifugation, fractions containing vector genomes were collected and pooled. The purified vectors were then concentrated using Amicon® Ultra-4 Centrifugal Filter Units (UFC810096). Each virus was washed three times with PBS at 4°C to remove residual CsCl. The resulting vector, and viral stocks were aliquoted and stored at −80°C. The detailed production protocol can be found in [81]. AAV titers were determined by SYBR Green qPCR against a standard curve and primer specific to the U6 promotor of the virus. U6/R1: 5’-gcctatttcccatgattcctt-3’. U6/L1: 5’-aaaactgcaaactacccaagaa-3’. The annealing temperature for the primers was 60°C.

### Cell Culture

The HEK293T/pBK1340 cell line was generated through transduction of HEK293T cells (ATCC® CRL3216™) with pLenti-pBK1340 and selected using puromycin. These cells express dGFP and luciferase (NLuc) downstream of a CMV promoter. The dGFP tag provides visual changes of gene repression whereas the luciferase provides higher spatial resolution identifying protein concentration changes over time. Maintenance cells were grown in DMEM (Gibco), supplemented with 10% fetal bovine serum (Gibco), penicillin/streptomycin 1% (Thermo Fisher Scientific), 2mM L-glutamine, 1% MEM NEAA (Gibco), and 1 mM sodium pyruvate (Gibco). To note: the cells were transduced with the MOI=0.1 to ensure 1 copy/cell following the selection. Before AAV transduction, the cells were seeded at 2.5 x 10^5^ cells per 12-well plate and cultured in DMEM with 2% FBS with no supplements. HEPG2 cells were maintained in DMEM (Gibco), supplemented with 10% fetal bovine serum (Gibco), penicillin/streptomycin 1% (Thermo Fisher Scientific), 2mM L-glutamine, 1% MEM NEAA (Gibco), and 1 mM sodium pyruvate (Gibco).

### AAV Transduction

At 50% confluency, the HEK293T/pBK1340 line was transduced with AAV/dCas9-repressor vectors. The following vectors were used: AAV/dCjCas9-HP1a +/-hinge region (fig.2b); AAV/dCjCas9-HP1b +/-hinge region (Fig.2b); AAV/dCjCas9-MBD1; AAV/dCjCas9-MBD2; AAV/dCjCas9-MBD3; AAV/dCjCas9-NIPP1; AAV/dCjCas9-MeCP2(TRD); AAV/dCjCas9-KRAB; AAV/dCjCas9-KRAB-MeCP2(TRD); AAV/dSaCas9-HP1a +/-hinge region (fig.2b); AAV/dSaCas9-HP1b +/-hinge region (fig.2b); AAV/dSaCas9-MBD1; AAV/dSaCas9-MBD2; AAV/dSaCas9-MBD3; AAV/dSaCas9-NIPP1; AAV/dSaCas9-MeCP2(TRD); AAV/dSaCas9-KRAB; AAV/dSaCas9-KRAB-MeCP2(TRD). The vectors were used with two different gRNAs (gRNA1 and gRNA2) (**Fig 2f**). Three different gRNA were used for targeting Pcsk9 promoter (gRNA1-3) (the sequences are listed above). The vectors were used at the MOIs=50,000 vg/cell. At 48 hours post transduction, the cells reached 100% confluency and were split to 40% confluency. Over the course of 21 or 28 days, the cells were passaged and harvested at 70% confluency to prevent epigenetic modifications caused by over confluency.

### Western Blotting

Expression levels of dGFP or Pcsk9 proteins in the stably transduced HEK293T cells or in HEPG2 cells were determined by western blotting with the GFP rabbit monoclonal antibody (ab138501, Abcam; 1:1,000) and with the Pcsk9 rabbit polyclonal antibody Catalog # 55206-1-AP from Thermo Fisher monoclonal antibody (mAb) β-actin (AM4302, Ambion; 1:5,000) for normalization. Cells were scraped from the dish and homogenized in 10× volume of 50 mM Tris-HCl (pH 7.5), 150 mM NaCl, 1% Nonidet P-40, in the presence of a protease and phosphatase inhibitor cocktail (Sigma, St. Louis, MO). Samples were sonicated 3 times for 15 s each cycle. Total protein concentrations were determined by the DC Protein Assay (Bio-Rad, Hercules, CA), and 25 μg of each sample was run on 12% Tris-glycine SDS-PAGE gels. Proteins were transferred to nitrocellulose membranes, and blots were blocked with 5% milk PBS Tween 20. Primary antibodies were incubated at 4°C overnight (Abcam, ab138501, 1:1,000; Thermo Fisher Scientific, AM4302, 1:5,000). Horseradish peroxidase-conjugated secondary antibodies were incubated for 1 hr at room temperature (Abcam; 1:10,000). Signal was detected with HyGLO Quick Spray (Denville Scientific) and immunoblots were imaged using ChemiDoc MP Imaging System (Bio-Rad). The densitometry was measured using ImageJ software, and α-synuclein expression was normalized to β-actin expression in the same lane. The experiment was repeated twice and represents two independent biological replicates.

### Luciferase Reporter Assay

Cells from each 12-well plate were first harvested and washed twice with 1xPBS before being resuspended in 200 µL of 1xPBS. 50 µL of the cell and 1xPBS mixture were transferred into a 96-well plate bottom white plate (Costar Cat#3922) and lysis buffer from Nano-Glo Luciferase Assay Kit (Promega Cat#N1120) was added directly to the plates following the manufacturer’s protocol. The data was obtained using a microplate spectrophotometer (Bio-Rad, Hercules, CA). Total protein concentration, which was determined by DC Protein Assay Set (Bio-Rad, Hercules, CA), was then used for data normalization.

### Genome DNA Extraction and AAV Integration Analysis

Genome DNA was extracted from HEK293T cells transduced with the dCas9-KRAB-MeCP2 viruses (first and second generation, −/+ Sp1-NF-kB, respectively or naïve AAV vector. The samples were harvested at the following time points: 2days, 7days, 14 days, 21 days, and 28 days post-transduction. The DNeasy Blood and Tissue Kit (QIAGEN) was used for the DNA isolation, per the manufacturer’s instructions. Next, the samples were digested with RNase A and DpnI overnight at 37°C. qPCR was used to quantify the level of AAV integration by replicating viral genome The following primers were used to amplify vector DNA: U6/R1: gcctatttcccatgattcctt; U6/L1: aaaactgcaaactacccaagaa; WPRE/R: actgtgtttgctgacgcaac; WPRE/L: agtcccggaaaggagctg. We used-beta-actin as a reference gene; Actin-F-5′-AATCTGCCACCACACCTTC-3′ and Actin-R-5′-GGGGTGTTGAAGGTCTCAAA-3′. iTaq Universal SYBR Green Supermix was used for the reactions (Bio-Rad). Real-time PCR was carried out using the iCycler iQ System (Bio-Rad), and the results were analyzed by iCycler software (Bio-Rad).

### Chromatin Immunoprecipitation (ChIP) Assay

This protocol was performed as described in [39] with slight modifications. HEK293T-reporter cells (see above), or HEPG2 cells were transduced with relevant viruses at an MOIs=50,000. Transduced cells were then cross-linked with 1% formaldehyde solution, (1% formaldehyde, 10 mM NaCl, 100 mM EDTA, 50 mM EGTA, 5 mM HEPES, pH 8.0) and quenched with 125 mM glycine. Cells were lysed with LB1 buffer (100 mM HEPES-KOH, pH 7.5, 280 mMNaCl, 2 mM EDTA, 20% glycerol, 1% Nonidet P-40, 0.5% Triton X-100), washed with LB2 buffer (400 mM NaCl, 2 mM EDTA, 1 mM EGTA, 20 mM Tris, pH 8.0), and resuspended in LB3 buffer (2 mM EDTA, 1 mM EGTA, 20 mM Tris, pH 8.0). Lysates were sonicated 12 times for 30 s, using a cell disruptor at output power 4 (Ultrasonic 350). Debris was precipitated at 14,000 rpm and 4 °C, and the sample was incubated with anti-histone H3, anti-acetylated H3 (SAB5700141 Millipore), anti-dimethylated H3-K4 (SAB5700160, Millipore); or anti-trimethylated H3-K9 (SAB5700163, Millipore) antibodies bound to protein A agarose beads (10001D, Thermo Fisher). Before incubation, a 1/50 input fraction was withdrawn. Beads were washed seven times with RIPA buffer (50 mM HEPES, pH 7.6, 1 mM EDTA, 0.7% DOC, 1% Nonidet P-40, 0.5 M LiCl) and once with TE buffer (50 mM Tris, pH 8.0, 10 mM EDTA). Beads were resuspended in elution buffer (50 mM Tris, pH 8.0, 10 mM EDTA, 1% SDS), incubated at 65 °C for 15 min with continuous shaking, and spun down. SN (bound fraction) and input fraction (withdrawn earlier) were incubated overnight at 65 °C to achieve reverse cross-linking. DNA was isolated using the phenol/chloroform protocol and resuspended in 50 uL double-distilled water. One microliter 1:4 diluted sample and 1 uL sample were amplified by real-time PCR. Primers used for PCR were as follows: Primers for GAPDH gene: Upper, 5’-TTCATCCAAGCGTGTAAGGG-3; lower, 5’-TGGTTCCCAGGACTGGACTGT-3’. Primers for beta-globin gene: Upper, 5’-CAGAGCCATCTATTGCTTAC-3’; lower, 5’-GCCTCACCACCAACTTCATC-3’. Primers for CMV promoter: Upper, 5’-GCAGTACATCTACGTATTAG-3’; lower, 5’-AGGTCAAAACAGCGTGGATG ‘3. Primers for Pcsk9 amplification were as follows: hPCSK9-qRT-R 5’-ccttcttcctggcttcctg-3’; hPCSK9-qRT-L 5’-gctctgggcaaagacagag-3’. The experiments were performed in triplicate. All statistical analysis was performed using GraphPad Prism 9. To determine statistical significance, Shapiro-Wilk tests were first used to evaluate the assumption of normality of the data. For normally distributed data Two-way ANOVA or t-test was used where appropriate, for non-normally distributed data Mann-Whitney test was used.

### Stereotaxic injections into the mouse hippocampus

Male and female C57BL/6 mice weighing 24–30g were obtained from Charles River Laboratories and investigated at the ages of 16 weeks or 32 weeks. Mice were keptunder standard conditions (21 °C, 12 h/12 h light-dark cycle) with food and water available *ad libitum* in their home cages. All animal protocols were approved by the Duke Institutional Animal Care and Use Committee. Under isoflurane anesthesia (0.5–2% isoflurane in O2) mice were injected bilaterally into the dorsal hippocampus (DH) via a Neuros syringe (Hamilton) at a rate of 150 nl/minute using the following stereotaxic coordinates (relative to bregma): 1.75 mm anterior, ± 1.5 mm lateral, and 1.55 mm ventral. Mice of 16 weeks and 32 weeks received combination of LV-dGFP-nLuc (LV-GFP) reporter and AAV/gRNA1-d*Sa*Cas9-KRAB-MeCP2(TRD) repressor vector (1 µl) in the left DH, and a combination of LV-GFP reporter and the control vector (no repressor, no gRNA) AAV/d*Sa*Cas (1 µl) in the right DH, providing a within animal control. Mice of 16 weeks received, AAV/gRNA_1 or 2_(*Apoe*)p-d*Sa*Cas9-KRAB-MeCP2(TRD) repressor vector (1 µl) in the right DH and the control vector AAV/d*Sa*Cas (1 µl) in the left DH. Animal welfare and weights were monitored daily for 14 days or 42 days.

### Immunohistochemistry and Imaging

At 14 days or 42 days post-injection, mice were transcardially perfused with 10% formalin and coronal slices (100 μm) containing the hippocampus were cut with a vibratome (VT1000S, Leica Microsystems). For immunohistochemistry, brain slices were immersed for 1 h at room temperature in PBS containing 0.2% Triton-X 100, 5% normal goat serum and 1x fish gelatin, then incubated with primary antibody for ApoE (1:800, ab183597, Abcam) at 4°C overnight on a rotator. The next day, the slices were rinsed 3 times in PBS and incubated in the corresponding secondary antibody (1:500, Invitrogen, thermo Fisher Scientific) at 4°C overnight on a rotator. Slices were then washed three times with PBS, mounted with VECTASHIELD + DAPI and cover-slipped. All slices were imaged under a 2x objective on a fluorescence microscope (Keyence BZ-X810). Images were taken of 3 slices per mouse and fluorescence intensity was analyzed in ImageJ.

### RNA Extraction and Expression Analysis

At 14 days or 42 days post-injection the left and right DH was dissected and immersed in an RNAlater™ solution (Thermo Fisher). Total RNA was extracted from DH samples using TRIzol reagent (Invitrogen) followed by purification with a RNeasy Mini Kit (Qiagen) according to the manufacturer’s protocol. RNA concentration was determined spectrophotometrically at 260 nm, while the quality of the purified RNA was determined by 260 nm/280 nm ratio. All of the RNA samples were of acceptable quality having ratios between 1.9 and 2.1. cDNA was synthesized using MultiScribe RT enzyme (Applied Biosystems) under the following conditions: 10 min at 25°C, 120 min at 37°C, 5 min at 85°C, and hold at 4°C.

Real-time PCR was used to quantify the level of GFP mRNA. Duplicates of each sample were assayed by relative quantitative real-time PCR using the ABI QuantStudio 7 to determine the level of the GFP mRNA relative to the mouse mRNAs for the housekeeping genes glyceraldehyde-3-phosphate dehydrogenase (mGapdh) and cyclophilin A (mPpia). ABI MGB probe and primer set assays were used to amplify the GFP cDNA; and the two RNA reference controls of the mouse endogenous, Gapdh (ID Mm99999915_g1, 109bp) and Ppia (ID Mm02342429_g1, 112bp) (Applied Biosystems). Each cDNA (10 ng) was amplified in duplicate in at least two independent runs (overall ≥ 4 repeats), using TaqMan Universal PCR master mix reagent (Applied Biosystems) and the following conditions: 2 min at 50°C, 10 min at 95°C, 40 cycles; 15 s at 95°C; and 1 min at 60°C. As a negative control for the specificity of the amplification, we used duplicates of no-templates (no-cDNA) in each plate, containing only probes and master mix. No amplification product was detected in control reactions. Data were analyzed with a threshold set in the linear range of amplification. The cycle number at which a sample crossed that threshold (Ct) was then used to determine fold difference, whereas the geometric mean of the two control genes served as a reference for normalization. Fold change was calculated as 2− ΔΔCt; where ΔCt=[Ct(target)-Ct (reference)] and ΔΔCt=[ΔCt(sample)]-[ ΔCt(calibrator)]. The calibrator was a dedicated RNA sample used in every plate for normalization within and across runs. The variation of the ΔCt values among the calibrator replicates was less than 10%.

### Statistical Analysis

For the cell culture experiments, the significance of the differences between no sgRNA and with sgRNA groups were analyzed statistically using student’s t test (Realstats Excel). Gene expression was measured via changes in luciferase concentration using a NanoLuc assay. A. A BCA Assay was also conducted to obtain the overall amino acid concentration and to normalize the data. The treated cells were then normalized using the control group of cells transfected/transduced with no sgRNA and no transcriptional repressor domain. Equations (1-2) demonstrate how the luciferase was normalized and quantified.

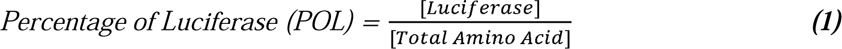

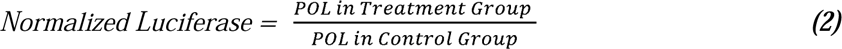

For the mouse experiments, all statistical analysis was performed using GraphPad Prism 9. To determine statistical significance, Shapiro-Wilk tests were first used to evaluate the assumption of normality of the data. For normally distributed data Two-way ANOVA or t-test was used where appropriate, for non-normally distributed data Mann-Whitney test was used.

## Disclosure

Drs. Chiba-Falek and Kantor are inventors of intellectual property related to this research and Duke University filed a patent application for the technology developed in this study. CLAIRIgene has an exclusive, worldwide option agreement from Duke for the related patent portfolio for all fields of use. Drs. Kantor and Chiba-Falek are Co-Founders at CLAIRIgene, LLC.

---

**Supplementary Figure 1:**
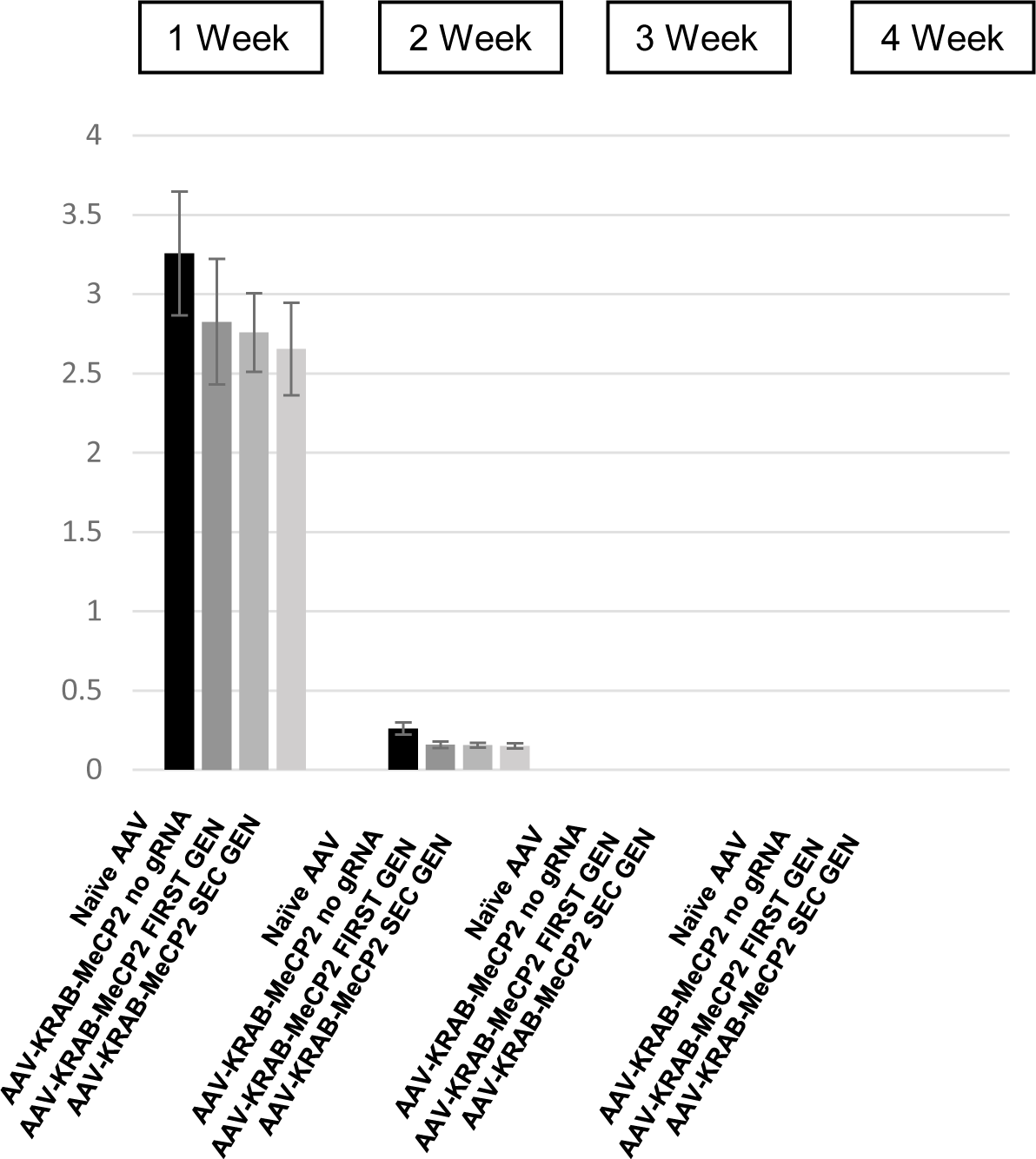

**Supplementary Figure 2:**
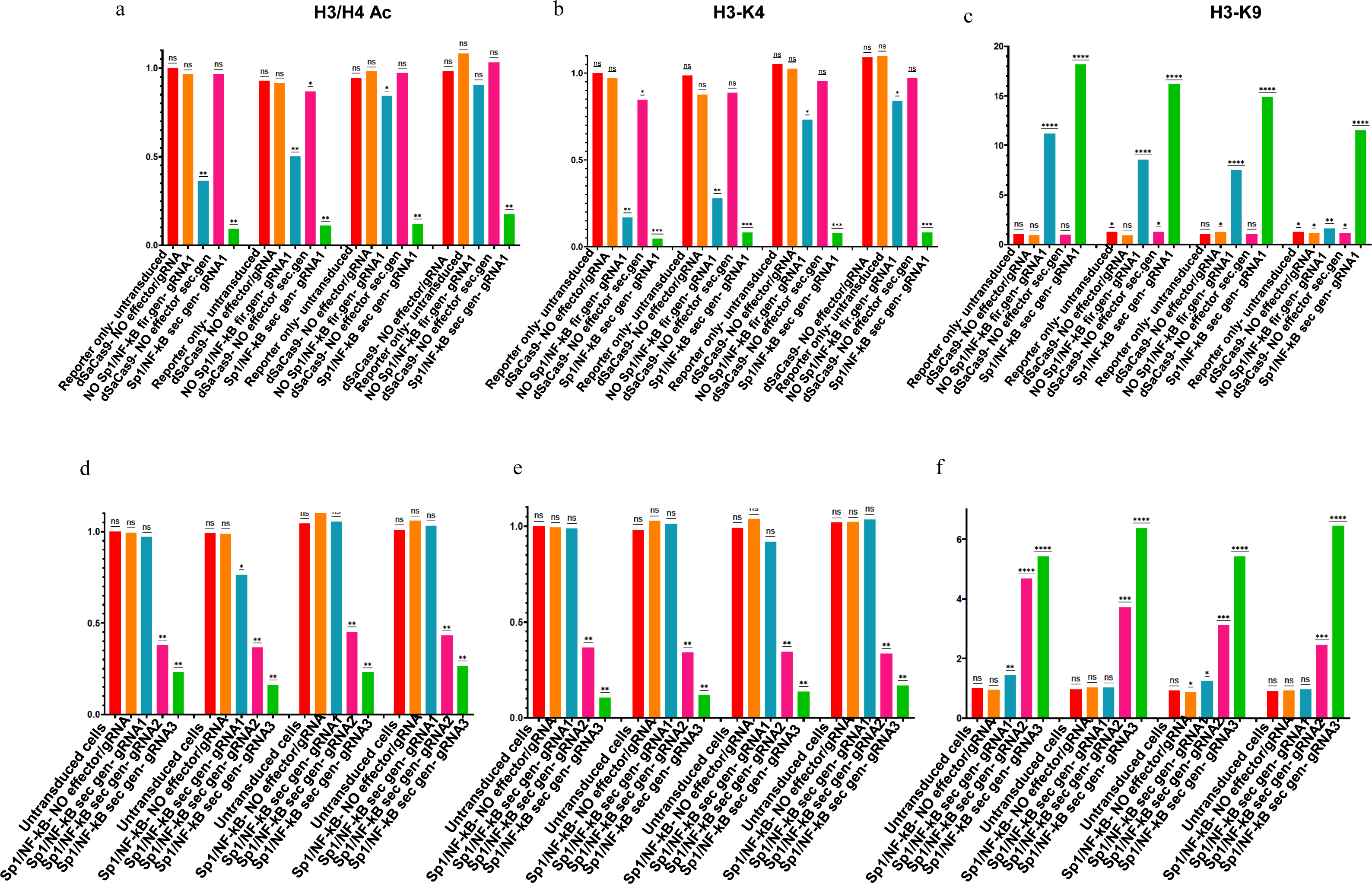

**Supplementary Figure 3:**
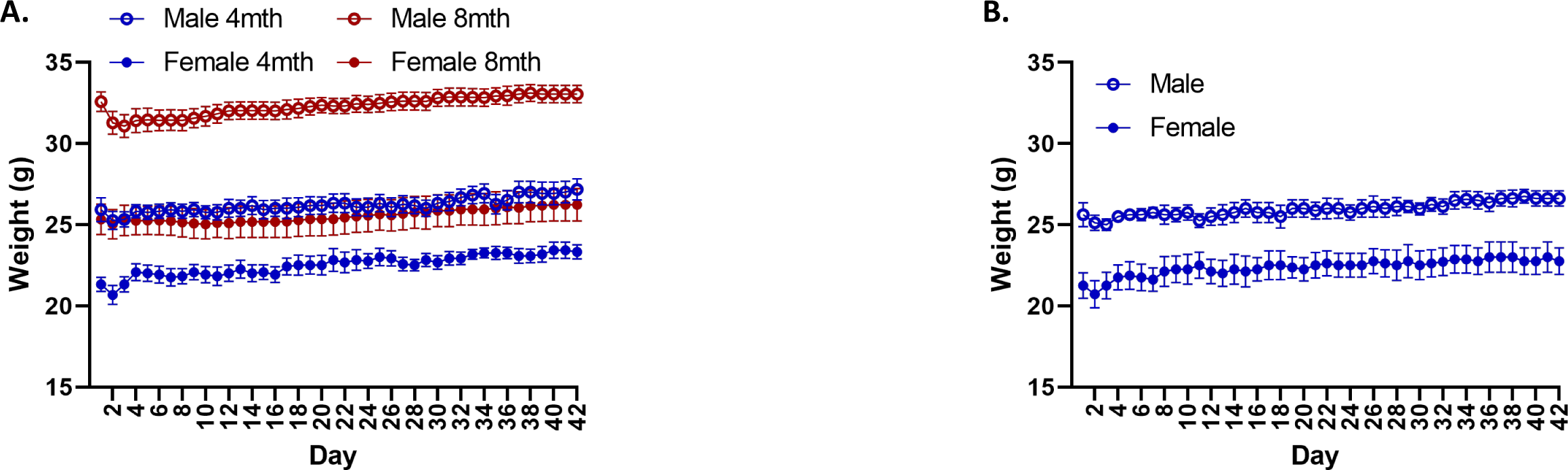
*In vivo* safety: There was no significant effect of **A.** The AAV-gRNA(GFP)/dCas-KRAB/MeCP2 vector, or **B.** The AAV-gRNA1(ApoE)/dCas-KRAB/MeCP2 and AAV-gRNA2(ApoE)/dCas-KRAB/MeCP2 vectors on mouse weights following injection. (16 week N=11, 32 week N=10. ApoE gRNA1 N=8 mice, ApoE gRNA2 N=8) Values represent mean ± SEM.

## References

1. Jinek, M., et al., A programmable dual-RNA-guided DNA endonuclease in adaptive bacterial immunity. Science, 2012. 337(6096): p. 816-21.

2. Cong, L., et al., Multiplex genome engineering using CRISPR/Cas systems. Science, 2013. 339(6121): p. 819-23.

3. Mali, P., et al., RNA-guided human genome engineering via Cas9. Science, 2013. 339(6121): p. 823-6.

4. Conti, A. and R. Di Micco, p53 activation: a checkpoint for precision genome editing? Genome Med, 2018. 10(1): p. 66.

5. Mandegar, M.A., et al., CRISPR Interference Efficiently Induces Specific and Reversible Gene Silencing in Human iPSCs. Cell Stem Cell, 2016. 18(4): p. 541–53.

6. Tsai, S.Q. and J.K. Joung, Defining and improving the genome-wide specificities of CRISPR-Cas9 nucleases. Nat Rev Genet, 2016. 17(5): p. 300–12.

7. Bikard, D., et al., Programmable repression and activation of bacterial gene expression using an engineered CRISPR-Cas system. Nucleic Acids Res, 2013. 41(15): p. 7429–37.

8. Gilbert, L.A., et al., CRISPR-mediated modular RNA-guided regulation of transcription in eukaryotes. Cell, 2013. 154(2): p. 442–51.

9. Maeder, M.L., et al., CRISPR RNA-guided activation of endogenous human genes. Nat Methods, 2013. 10(10): p. 977–9.

10. Wu, X., et al., Genome-wide binding of the CRISPR endonuclease Cas9 in mammalian cells. Nat Biotechnol, 2014. 32(7): p. 670–6.

11. Qi, L.S., et al., Repurposing CRISPR as an RNA-guided platform for sequence-specific control of gene expression. Cell, 2013. 152(5): p. 1173–83.

12. Tanenbaum, M.E., et al., A protein-tagging system for signal amplification in gene expression and fluorescence imaging. Cell, 2014. 159(3): p. 635–46.

13. Larson, M.H., et al., CRISPR interference (CRISPRi) for sequence-specific control of gene expression. Nat Protoc, 2013. 8(11): p. 2180–96.

14. Konermann, S., et al., Genome-scale transcriptional activation by an engineered CRISPR-Cas9 complex. Nature, 2015. 517(7536): p. 583-8.

15. Rittiner, J.E., et al., Gene-Editing Technologies Paired With Viral Vectors for Translational Research Into Neurodegenerative Diseases. Front Mol Neurosci, 2020. 13: p. 148.

16. Vojta, A., et al., Repurposing the CRISPR-Cas9 system for targeted DNA methylation. Nucleic Acids Res, 2016. 44(12): p. 5615–28.

17. Liu, X.S., et al., Editing DNA Methylation in the Mammalian Genome. Cell, 2016. 167(1): p. 233–247 e17.

18. Liu, X.S., et al., Rescue of Fragile X Syndrome Neurons by DNA Methylation Editing of the FMR1 Gene. Cell, 2018. 172(5): p. 979–992 e6.

19. Kantor, B., et al., Downregulation of SNCA Expression by Targeted Editing of DNA Methylation: A Potential Strategy for Precision Therapy in PD. Mol Ther, 2018. 26(11): p. 2638–2649.

20. Chen, V., et al., The mechanistic role of alpha-synuclein in the nucleus: impaired nuclear function caused by familial Parkinson’s disease SNCA mutations. Hum Mol Genet, 2020. 29(18): p. 3107–3121.

21. Kantor, B., et al., Methods for gene transfer to the central nervous system. Adv Genet, 2014. 87: p. 125–97.

22. Ortinski, P.I., et al., Integrase-Deficient Lentiviral Vector as an All-in-One Platform for Highly Efficient CRISPR/Cas9-Mediated Gene Editing. Mol Ther Methods Clin Dev, 2017. 5: p. 153–164.

23. Kantor, B., et al., Clinical applications involving CNS gene transfer. Adv Genet, 2014. 87: p. 71–124.

24. Rittiner, J., et al., Therapeutic modulation of gene expression in the disease state: Treatment strategies and approaches for the development of next-generation of the epigenetic drugs. Front Bioeng Biotechnol, 2022. 10: p. 1035543.

25. Ran, F.A., et al., In vivo genome editing using Staphylococcus aureus Cas9. Nature, 2015. 520(7546): p. 186-91.

26. Kim, E., et al., In vivo genome editing with a small Cas9 orthologue derived from Campylobacter jejuni. Nat Commun, 2017. 8: p. 14500.

27. Burstein, D., et al., New CRISPR-Cas systems from uncultivated microbes. Nature, 2017. 542(7640): p. 237-241.

28. Xu, X., et al., Engineered miniature CRISPR-Cas system for mammalian genome regulation and editing. Mol Cell, 2021. 81(20): p. 4333–4345 e4.

29. Thakore, P.I., et al., RNA-guided transcriptional silencing in vivo with S. aureus CRISPR-Cas9 repressors. Nat Commun, 2018. 9(1): p. 1674.

30. Chavez, A., et al., Comparison of Cas9 activators in multiple species. Nat Methods, 2016. 13(7): p. 563–567.

31. Nakamura, M., et al., CRISPR technologies for precise epigenome editing. Nat Cell Biol, 2021. 23(1): p. 11–22.

32. Yeo, N.C., et al., An enhanced CRISPR repressor for targeted mammalian gene regulation. Nat Methods, 2018. 15(8): p. 611–616.

33. Duke, C.G., et al., An Improved CRISPR/dCas9 Interference Tool for Neuronal Gene Suppression. Front Genome Ed, 2020. 2: p. 9.

34. Kantor, B., et al., Notable reduction in illegitimate integration mediated by a PPT-deleted, nonintegrating lentiviral vector. Mol Ther, 2011. 19(3): p. 547–56.

35. Poon, A.P., H. Gu, and B. Roizman, ICP0 and the US3 protein kinase of herpes simplex virus 1 independently block histone deacetylation to enable gene expression. Proc Natl Acad Sci U S A, 2006. 103(26): p. 9993–8.

36. Nevels, M., C. Paulus, and T. Shenk, Human cytomegalovirus immediate-early 1 protein facilitates viral replication by antagonizing histone deacetylation. Proc Natl Acad Sci U S A, 2004. 101(49): p. 17234–9.

37. Richart, A.N., et al., Characterization of chromoshadow domain-mediated binding of heterochromatin protein 1alpha (HP1alpha) to histone H3. J Biol Chem, 2012. 287(22): p. 18730–7.

38. Monahan, P.E., et al., Proteasome inhibitors enhance gene delivery by AAV virus vectors expressing large genomes in hemophilia mouse and dog models: a strategy for broad clinical application. Mol Ther, 2010. 18(11): p. 1907–16.

39. Kantor, B., et al., Epigenetic activation of unintegrated HIV-1 genomes by gut-associated short chain fatty acids and its implications for HIV infection. Proc Natl Acad Sci U S A, 2009. 106(44): p. 18786–91.

40. Pereira, D.J. and N. Muzyczka, The cellular transcription factor SP1 and an unknown cellular protein are required to mediate Rep protein activation of the adeno-associated virus p19 promoter. J Virol, 1997. 71(3): p. 1747–56.

41. Van Lint, C., et al., A transcriptional regulatory element is associated with a nuclease-hypersensitive site in the pol gene of human immunodeficiency virus type 1. J Virol, 1994. 68(4): p. 2632–48.

42. Goffin, V., et al., Transcription factor binding sites in the pol gene intragenic regulatory region of HIV-1 are important for virus infectivity. Nucleic Acids Res, 2005. 33(13): p. 4285–310.

43. Lee, D.H., et al., Comparison of three heterochromatin protein 1 homologs in Drosophila. J Cell Sci, 2019. 132(3).

44. Smothers, J.F. and S. Henikoff, The hinge and chromo shadow domain impart distinct targeting of HP1-like proteins. Mol Cell Biol, 2001. 21(7): p. 2555–69.

45. Razin, A. and B. Kantor, DNA methylation in epigenetic control of gene expression. Prog Mol Subcell Biol, 2005. 38: p. 151–67.

46. Dong, W. and B. Kantor, Lentiviral Vectors for Delivery of Gene-Editing Systems Based on CRISPR/Cas: Current State and Perspectives. Viruses, 2021. 13(7).

47. Policarpi, C., J. Dabin, and J.A. Hackett, Epigenetic editing: Dissecting chromatin function in context. Bioessays, 2021. 43(5): p. e2000316.

48. Abifadel, M., et al., Mutations in PCSK9 cause autosomal dominant hypercholesterolemia. Nat Genet, 2003. 34(2): p. 154–6.

49. Cohen, J.C., et al., Sequence variations in PCSK9, low LDL, and protection against coronary heart disease. N Engl J Med, 2006. 354(12): p. 1264–72.

50. Vercruysse, P., et al., Hypothalamic Alterations in Neurodegenerative Diseases and Their Relation to Abnormal Energy Metabolism. Front Mol Neurosci, 2018. 11: p. 2.

51. Yang, A., B. Kantor, and O. Chiba-Falek, APOE: The New Frontier in the Development of a Therapeutic Target towards Precision Medicine in Late-Onset Alzheimer’s. Int J Mol Sci, 2021. 22(3).

52. Nan, X., et al., Transcriptional repression by the methyl-CpG-binding protein MeCP2 involves a histone deacetylase complex. Nature, 1998. 393(6683): p. 386-9.

53. Nan, X., F.J. Campoy, and A. Bird, MeCP2 is a transcriptional repressor with abundant binding sites in genomic chromatin. Cell, 1997. 88(4): p. 471–81.

54. Hendrich, B. and A. Bird, Identification and characterization of a family of mammalian methyl-CpG binding proteins. Mol Cell Biol, 1998. 18(11): p. 6538–47.

55. Kantor, B., et al., Expression and localization of components of the histone deacetylases multiprotein repressory complexes in the mouse preimplantation embryo. Gene Expr Patterns, 2003. 3(6): p. 697–702.

56. Chew, W.L., et al., A multifunctional AAV-CRISPR-Cas9 and its host response. Nat Methods, 2016. 13(10): p. 868–74.

57. Moreno, A.M., et al., In Situ Gene Therapy via AAV-CRISPR-Cas9-Mediated Targeted Gene Regulation. Mol Ther, 2020. 28(8): p. 1931.

58. Liao, H.K., et al., In Vivo Target Gene Activation via CRISPR/Cas9-Mediated Trans-epigenetic Modulation. Cell, 2017. 171(7): p. 1495–1507 e15.

59. Jost, M., et al., Titrating gene expression using libraries of systematically attenuated CRISPR guide RNAs. Nat Biotechnol, 2020. 38(3): p. 355–364.

60. Amabile, A., et al., Inheritable Silencing of Endogenous Genes by Hit-and-Run Targeted Epigenetic Editing. Cell, 2016. 167(1): p. 219–232 e14.

61. Van, M.V., T. Fujimori, and L. Bintu, Nanobody-mediated control of gene expression and epigenetic memory. Nat Commun, 2021. 12(1): p. 537.

62. Mlambo, T., et al., Designer epigenome modifiers enable robust and sustained gene silencing in clinically relevant human cells. Nucleic Acids Res, 2018. 46(9): p. 4456–4468.

63. Vijayraghavan, S. and B. Kantor, A Protocol for the Production of Integrase-deficient Lentiviral Vectors for CRISPR/Cas9-mediated Gene Knockout in Dividing Cells. J Vis Exp, 2017(130).

64. Sun, Z., B. Kantor, and O. Chiba-Falek, Neuronal-type-specific epigenome editing to decrease SNCA expression: Implications for precision medicine in synucleinopathies. Mol Ther Nucleic Acids, 2024. 35(1): p. 102084.

65. Babcock, K.R., et al., Adult Hippocampal Neurogenesis in Aging and Alzheimer’s Disease. Stem Cell Reports, 2021. 16(4): p. 681–693.

66. Li, Y.J., et al., Identification of novel genes for age-at-onset of Alzheimer’s disease by combining quantitative and survival trait analyses. Alzheimers Dement, 2023.

67. Coon, K.D., et al., A high-density whole-genome association study reveals that APOE is the major susceptibility gene for sporadic late-onset Alzheimer’s disease. Journal of Clinical Psychiatry, 2007. 68: p. 613–618.

68. Grupe, A., et al., Evidence for novel susceptibility genes for late-onset Alzheimer’s disease from a genome-wide association study of putative functional variants. Human Molecular Genetics, 2007. 16: p. 865–873.

69. Ernoult, E., E. Gamelin, and C. Guette, Improved proteome coverage by using iTRAQ labelling and peptide OFFGEL fractionation. Proteome Sci, 2008. 6: p. 27.

70. Abraham, R., et al., A genome-wide association study for late-onset Alzheimer’s disease using DNA pooling. BMC Medical Genomics, 2008. 1: p. 44.

71. Bertram, L., et al., Genome-wide association analysis reveals putative Alzheimer’s disease susceptibility loci in addition to APOE. American Journal of Human Genetics, 2008. 83: p. 623–632.

72. Beecham, G.W., et al., Genome-wide association study implicates a chromosome 12 risk locus for late-onset Alzheimer disease. American Journal of Human Genetics, 2009. 84: p. 35–43.

73. Carrasquillo, M.M., et al., Genetic variation in PCDH11X is associated with susceptibility to late-onset Alzheimer’s disease. Nature Genetics, 2009. 41: p. 192–198.

74. Harold, D., et al., Genome-wide association study identifies variants at CLU and PICALM associated with Alzheimer’s disease. Nature Genetics, 2009. 41: p. 1088–1093.

75. Lambert, J.C., et al., Genome-wide association study identifies variants at CLU and CR1 associated with Alzheimer’s disease. Nature Genetics, 2009. 41: p. 1094–1099.

76. Potkin, S.G., et al., Hippocampal atrophy as a quantitative trait in a genome-wide association study identifying novel susceptibility genes for Alzheimer’s disease. PLoS One 4, 2009. 4: p. e6501.

77. Heinzen, E.L., et al., Genome-wide scan of copy number variation in late-onset Alzheimer’s disease. Journal of Alzheimer’s Disease. Journal of Alzheimer’s Disease, 2010. 19: p. 69–77.

78. Seshadri, S., et al., Genome-wide analysis of genetic loci associated with Alzheimer disease. JAMA, 2010. 303: p. 1832–1840.

79. Kamboh, M.I., et al., Genome-wide association analysis of age-of-onset in Alzheimer’s disease. Molecular Psychiatry, 2012. 17: p. 1340–1346.

80. Gottschalk, W.K., et al., The Role of Upregulated APOE in Alzheimer’s Disease Etiology. J Alzheimers Dis Parkinsonism, 2016. 6(1).

81. Grieger, J.C., V.W. Choi, and R.J. Samulski, Production and characterization of adeno-associated viral vectors. Nat Protoc, 2006. 1(3): p. 1412–28.

